# Estimation of demography and mutation rates from one million haploid genomes

**DOI:** 10.1101/2024.09.18.613708

**Authors:** Joshua G. Schraiber, Jeffrey P. Spence, Michael D. Edge

**Affiliations:** Department of Quantitative and Computational Biology, Univeristy of Southern California, USA; Department of Genetics, Stanford University, USA

## Abstract

As genetic sequencing costs have plummeted, datasets with sizes previously un-thinkable have begun to appear. Such datasets present new opportunities to learn about evolutionary history, particularly via rare alleles that record the very recent past. However, beyond the computational challenges inherent in the analysis of many large-scale datasets, large population-genetic datasets present theoretical problems. In particular, the majority of population-genetic tools require the assumption that each mutant allele in the sample is the result of a single mutation (the “infinite sites” assumption), which is violated in large samples. Here, we present DR EVIL, a method for estimating mutation rates and recent demographic history from very large samples. DR EVIL avoids the infinite-sites assumption by using a diffusion approximation to a branching-process model with recurrent mutation. The branching-process approach limits the method to rare alleles, but, along with recent results, renders tractable likelihoods with recurrent mutation. We show that DR EVIL performs well in simulations and apply it to rare-variant data from a million haploid samples, identifying a signal of mutation-rate heterogeneity within commonly analyzed classes and predicting that in modern sample sizes, most rare variants at sites with high mutation rates represent the descendants of multiple mutation events.

## 1 Introduction

We now have datasets with genome or exome sequences from hundreds of thousands of individuals [1–8]. Large genomic datasets have the potential to shed light on the evolutionary forces that shape genetic variation by enabling a high-resolution view of recent demographic history [9–12], mutation rates [1, 2, 4, 13–17], and natural selection [4, 18–20]. Understanding these evolutionary forces is crucial for the interpretation of whole-genome sequencing data.

From a population-genetic perspective, the key advantage of ultra-large sequencing datasets is the information they provide about rare variants. Because the age of a variant is correlated with its frequency [21–25], rare variants largely represent mutations that arose recently, and the frequencies of rare variants are informative about recent population history. This is especially relevant in humans, where recent explosive population growth has resulted in a large excess of rare variants [9].

In addition, rare variants can provide substantial power to estimate mutation rates. Although average mutation rates can be estimated from *de novo* mutations in trios or large pedigrees [26–28], there is substantial mutation-rate variation throughout the genome [13, 17, 29–31]. Many genomic features, such as flanking nucleotide sequence [13, 32], methylation level [32], and replication timing [31] are known to affect mutation rate. The many rare variants found in samples of hundreds of thousands of haplotypes provide a large pool of data from which to estimate mutation rates that depend on these features [17, 30]. Moreover, because the full set of factors that affect mutation rates is unknown, there may be residual mutation-rate heterogeneity that can be identified in large datasets, providing clues to help discover new factors that influence mutation rates.

Natural selection prevents most deleterious variants from rising to high frequency. Thus, rare variants are enriched for deleterious mutations. This fact is commonly invoked in criteria to predict variant pathogenicity [33–37] or in efforts to enrich for variants that are more likely to play a causal role in complex traits and disease [7, 38–40]. However, the interaction of mutation, selection, and demography means that simple rules for identifying putatively pathogenic variants, such as allele-frequency cutoffs, are not robust to variation in mutation rate [31]. Moreover, recent work showed that the distribution of allele frequencies, not merely the presence or absence of alleles, contributes substantially to improving estimates of selection [19]. It is therefore important to build models of rare variation that can account for natural selection and recurrent mutation to enhance our knowledge of natural selection in the human genome.

Despite the importance of rare variation in large datasets, most current populationgenetic methods are unable to access the information contained in such variants. One of the key assumptions undergirding many population-genetic methods is that all copies of a given allele share a single mutational origin, known as the infinite-sites assumption [41–43]. In contrast, recurrent mutation, in which variants of a given type have multiple mutational origins, is detectable even in datasets with only tens of thousands of individuals [1, 28, 44]. In addition, though there exist fast algorithms for computing the likelihood of genetic data in the absence of natural selection [45–48], computing likelihoods for variants subject to natural selection remains challenging [47, 49–51]. Although simulation-based approaches, such as approximate Bayesian computation [52] or supervised machine learning using simulations [53] can circumvent some of these limitations, the results of these approaches can be difficult to interpret in terms of the features of the input data, and the computational burden associated with simulation can make exploration of different models infeasible. Some recent approaches have made progress modeling rare variants subject to recurrent mutation [54] and selection [55, 56], clarifying how recurrent mutation and rapid population growth shape the distribution of allele frequencies. However, they are not geared toward estimation of mutation rates and demography.

To enable population-genetic inference from rare variation in ultra-large sequencing datasets, we present DR EVIL (<u>D</u>iffusion for <u>R</u> are <u>E</u>lements in <u>V</u>ariation Inventories that are <u>L</u>arge), a method for inference of recent demography and mutation rates using rare variants appearing in very large samples. The core of our method is a rare-variant approximation of the usual diffusion approximation used in population genetics that incorporates both recurrent mutation and selection. We show that the resulting process falls into a model class for which a solution was recently found, and we use the solution to arrive at approximate likelihoods for counts of rare alleles, which can then be optimized to estimate mutation and demographic parameters.

We compare the performance of our approach for mutation-rate estimation with standard methods in the literature and show that we improve estimation accuracy and are able to correct for the presence of mutation-rate heterogeneity. Furthermore, we demonstrate the importance of correctly accounting for recurrent mutation when estimating recent demographic history.

We then apply our method to one million samples from gnomAD, providing refined estimates of demographic history and mutation rates compared with the state of the art, including the detection of mutation-rate heterogeneity that remains after accounting for methylation status and trinucleotide context. Finally, we demonstrate the importance of accurate modeling of rare variation by exploring the impact of natural selection on the probability of observed allele counts as a function of mutation rate and the number of distinct mutations to which the copies of a rare allele in a very large sample trace their origin.

### 2 Results

## 2.1 Approximate sampling formula for rare alleles

We use a standard Wright–Fisher model of allele-frequency dynamics in a population of time-varying size. Specifically, we assume mutations arise with rate *µ* per site per generation, the fitness of the heterozygote is 1 + *hs*, and that the effective population size at time *t* is given by a function *N* (*t*). Importantly, we do not make the infinite-sites assumption that each site is only mutated once. Instead, we allow for recurrent mutation.

To model the site-frequency spectrum of a large sample, we require the probabilities *p_n,k_* that an allele in a sample of *n* haploid genomes will be observed *k* times. To obtain these probabilities for rare alleles (i.e. those for which *k ≪ n*), we make two approximations in addition to the standard Wright–Fisher diffusion approximation. First, we approximate the typical Wright–Fisher diffusion using Feller’s continuous-state branching process [57]. Branching-process approximations to the dynamics of rare alleles have a long history in population genetics [58, 59]. Intuitively, the branching process assumes that the allele is sufficiently rare that it is only found in heterozygous individuals, and additionally that heterozygous individuals are rare enough that they are unlikely to mate with other heterozygotes. Thus, they can be modeled as expanding in an unconstrained population (Figure 1, top left). Next, we approximate the binomial sampling of alleles in a finite sample with Poisson sampling [60]. Here, we rely on the fact that a binomial sample with a small probability of success in a large number of trials is well approximated by a Poisson distribution. Defining the relative population size *ρ*(*t*) = *N* (*t*)*/N*_0_ for some arbitrary *N*_0_, the population-scaled mutation rate *θ* = 4*N*_0_*µ*, and the population-scaled heterozygous selection coefficient *γ* = 4*N*_0_*hs*, we show in the Supplementary Material that the sampling 1probabilities satisfy a system of differential equations,

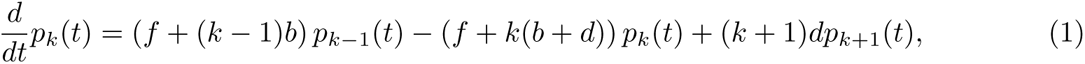

where 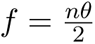 is an immigration rate, 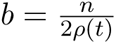is a birth rate, and 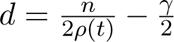 is a death rate. The formula (1) can then be recognized as an inhomogeneous linear birth-death process with immigration. Although birth-death processes with immigration have been studied for many years [61], to our knowledge the first analytic solution with inhomogeneous coefficients was presented recently [62]. In the Supplementary Material, we show that this solution can be written as

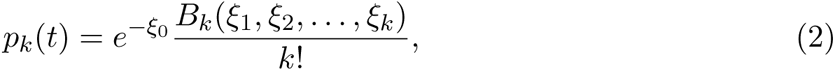

where *B_k_*(*x*_1_*, . . . , x_k_*) is a complete Bell polynomial [63, 64] whose arguments are given by

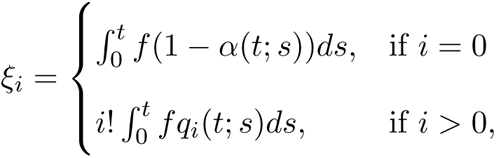

where *α*(*t*; *s*) is the probability that a birth-death process (without immigration) started with one copy at time *s* is not extinct at time *t* and *q_i_*(*t*; *s*) is the probability that a birth-death process (without immigration) started at one copy at time *s* is at *i* copies at time *t*. Explicit formulas for *α*(*t*; *s*) and *q_i_*(*t*; *s*) are given in the Supplementary Material. This formula has an interpretation as adding birth-death genealogies arising from each mutational origin (Figure 1, bottom left), and is explored further in the Supplementary Material. We also describe a reparameterization used to improve computational efficiency and numerical stability in the Supplementary Material. In simulations, we find that our analytic model matches the simulated site-frequency spectrum, whereas classic approaches using the infinite-sites model fail as sample sizes become larger (Figure 1, right side).

**Figure 1:**
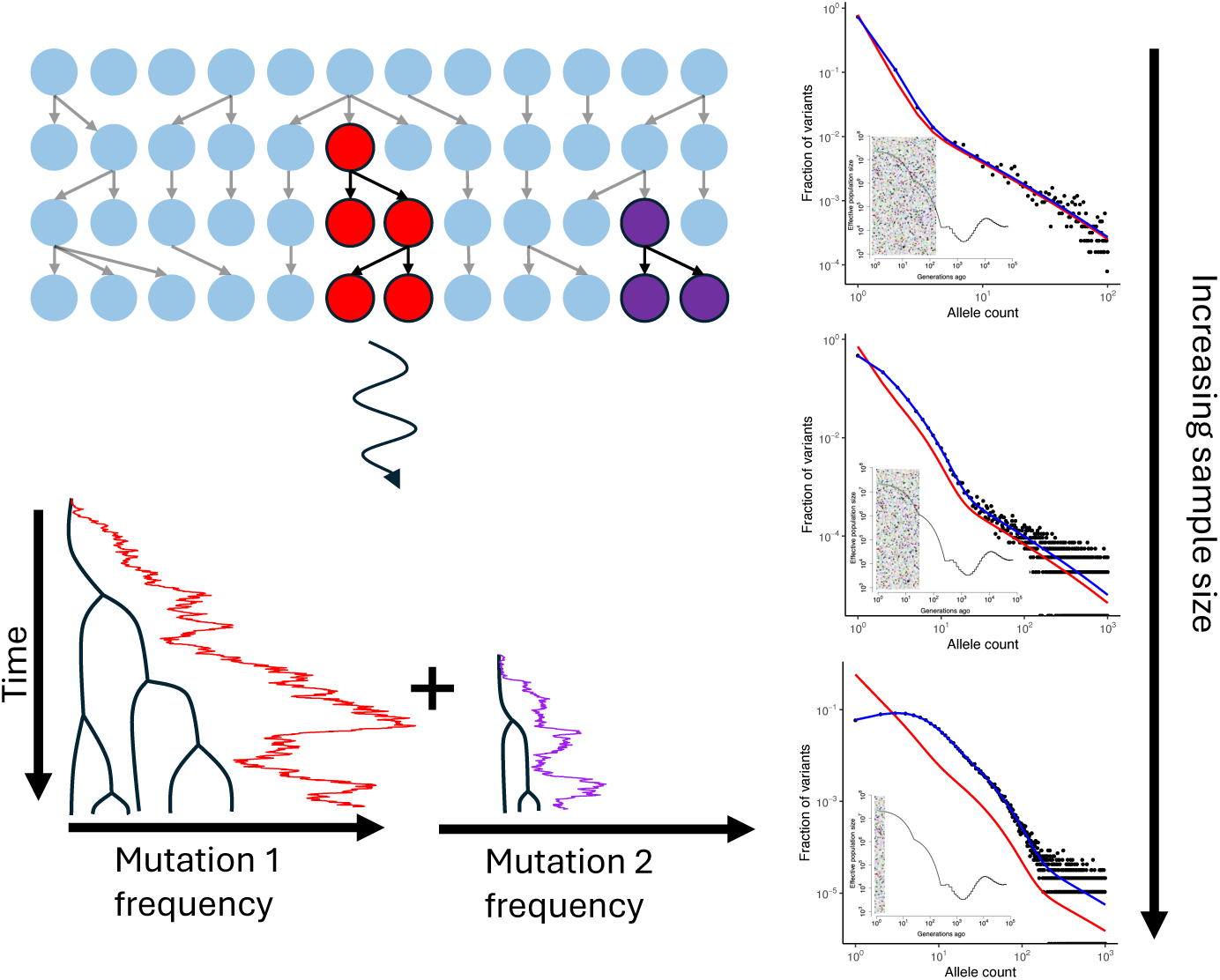
Modeling rare variation and recurrent mutation. On the left side, we consider a discrete population model in which a rare variant expands into a population that is dominated by another allele (indicated in blue). Two lineages of the rare variant arise, one shown in red and one shown in purple. We make a diffusion approximation to this process, as indicated by the curved arrow. In the diffusion approximation, we find that the genealogy of a sample of rare alleles is modeled by a birth-death process, and that the sampling distribution can be obtained by adding the sampling distribution of the two different origins. On the right side, we demonstrate how increasing sample sizes causes simulated data (shown as filled circles) to depart from an infinite-sites model (shown in red) and that our method (shown in blue), accurately matches the site frequency spectrum by modeling recurrent mutation. In each inset, we qualitatively indicate that as sample sizes become increasingly large, we are able to reveal more recent demographic history.

## 2.2 Maximum-likelihood inference of demography and mutation rates

In the Methods, we describe how to use the sampling formula (2) as part of a maximum-likelihood estimation procedure for mutation rates and demographic history. Here, we assess the performance of our method and compare it with other approaches.

### 2.2.1 Estimation of mutation rates

To examine the performance of our estimation procedure, we first explore the estimation of mutation rates when the demographic model is fixed. Although our method allows maximum-likelihood estimation of mutation rates, an alternative method-of-moments procedure is suggested by the solution in equation (2). In empirical data, the probability that a site is not variable, *p*_0_, can be estimated as one minus the fraction of segregating in the sample,

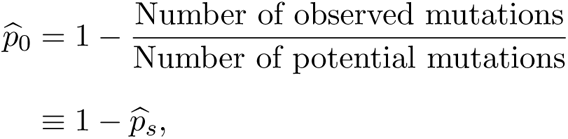

where we use 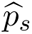 to indicate proportion of segregating sites in the sample. Then, noting that the immigration rate 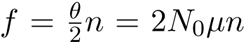, a method-of-moments estimator of the mutation

rate can be computed,

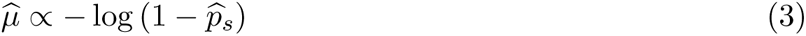

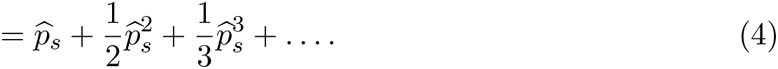

This method-of-moments estimator is implicit in [54] and used by [17]. The second line follows from a Taylor expansion of the logarithm, and the constant of proportionality can be determined by, for example, constraining the average mutation rate in the genome [2, 4, 16]. The first-order term, 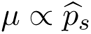, corresponds to an infinite-sites estimator of the mutation rate, and is commonly used to estimate mutation rates from population-genetic data [2, 4, 37]. However, it is clear from equation (4) that the first-order approximation is poor when the fraction of segregating sites is appreciable, underestimating the mutation rate. The full method-of-moments estimator also fails when a mutational context is fully saturated: in that case, 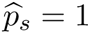 and the mutation-rate estimate is infinite. For this reason, mutation-rate estimates in high mutation-rate contexts are typically made by down-sampling data [2, 4], discarding information.

With DR EVIL, we use maximum likelihood to estimate the mutation rate. Maximum likelihood has the advantage of using allele-frequency information in addition to presence/absence information. When mutation rates are low, the mutation rate simply scales the frequency spectrum, and thus allele frequencies provide little information above presence/absence. However, when there are recurrent mutations, the shape of the site-frequency spectrum is influenced by the mutation rate (Figure 2A). Particularly of note is that as the mutation rate increases, the site frequency spectrum becomes non-monotonic and non-convex; under the infinite-sites model, the frequency spectrum must be decreasing and convex in the absence of population structure [65].

**Figure 2:**
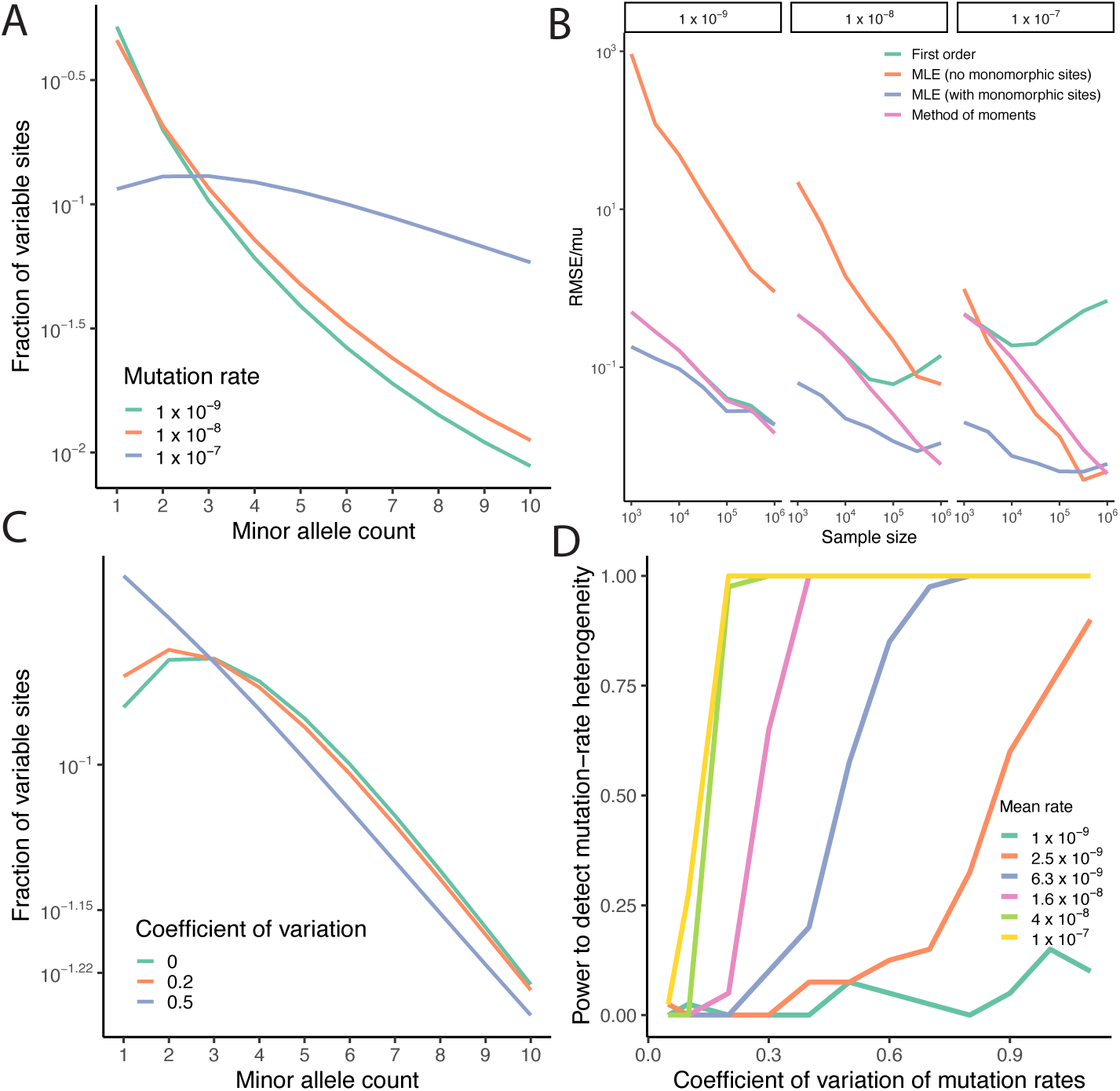
Mutation-rate estimation from site-frequency spectra with recurrent mutation. A) Three example site-frequency spectra, simulated with the demography from **Agarwal et al.** and three mutation rates, indicated by color. B) Performance of different methods of mutation-rate estimation as sample size changes. The horizontal axis shows the sample size and the vertical axis the relative root mean squared error. Different methods are indicated with different colors. Each panel shows the results of a simulation with the mutation rate indicated at the top of the panel. C) The impact of mutation-rate heterogeneity on the site-frequency spectrum. Each SFS has an average mutation rate of 10*^−^*^7^, but mutation rates are distributed according to a gamma distribution with the indicated coefficient of variation. D) Power to detect mutation-rate heterogeneity. The horizontal axis shows the coefficient of variation of the distribution of mutation rates, and the vertical axis shows the power to infer mutation-rate heterogeneity. Each line represents a different mutation rate. gamma distribution of mutation rates.

One downside of existing approaches and the DR EVIL MLE is that they require an accurate estimate of the mutational target size. However, due to the subtleties of read mapping and genotype calling, assessing which sites could have potentially been called as variable is surprisingly difficult [66, 67]. Because the shape of the site-frequency spectrum is influenced by the mutation rate, it provides information to estimate mutation rates without an estimate of the mutational target size, providing a method for mutation-rate estimation that is independent of some technical challenges. Thus, we also explored the possibility of estimating mutation rates without target sizes.

To test these different methods for inferring mutation rates, we simulated data under a recently proposed demographic model [18] while varying sample size and mutation rate. Figure 2B shows that the maximum-likelihood estimator of the mutation rate is consistently more accurate than any alternative; Supplementary Figure 2 shows that the other approaches are biased downward. As expected, for small mutation rates and sample sizes, the method-of-moments and first-order estimator perform equivalently. However, as the sample size increases, the method-of-moments and maximum-likelihood estimators become increasingly accurate, but the first-order estimator begins to increase in relative root mean squared error; this is because it does not properly account for recurrent mutation. Moreover, though the maximum-likelihood estimator without a target size performs poorly for low mutation rates and small sample sizes, it becomes comparable to the full maximum-likelihood estimator for high mutation rates at large sample sizes.

Finally, although it is common practice in human genetics to stratify variants by factors that influence their mutation rate (for example, trinucleotide context and methylation level), there is substantial evidence for residual mutation-rate variation [13, 17, 30]. To model mutation-rate variation that is not captured by the chosen stratification strategy, we incorporated a gamma distribution of mutation rates (Supplementary Material). We emphasize that we chose the gamma distribution for analytical convenience and flexibility and do not claim a biological motivation. Figure 2C shows that the coefficient of variation, which measures the amount mutation-rate heterogeneity, has a substantial impact on the shape of the frequency spectrum, resulting in an excess of low-frequency variants. To test the ability to detect residual mutation-rate variation, we simulated data under a gamma distribution of mutation rates. Figure 2D shows statistical power as a function of the coefficient of variation and the mutation rate. Unsurprisingly, there is more power to detect mutation-rate heterogeneity for higher average mutation rates and larger coefficients of variation. Supplementary Figure 3 shows that not accounting for mutation-rate heterogeneity results in a biased estimate of the average mutation rate. However, when we reject the null hypothesis of a single mutation rate, we accurately estimate the mean mutation rate and coefficient of variation.

### 2.2.2 Joint estimation of demography and mutation

In most applications, the exact demography is unknown and must also be estimated. This is particularly true when working with very large datasets: a key advantage of using extremely large sample sizes is that they reveal more recent demographic history because they contain rare variants that arose recently. However, in large samples, recurrent mutation distorts the site-frequency spectrum compared with what would be expected under an infinite-sites model, as seen in Figure 1. Therefore, inferences about demographic history made using tools that rely on the infinite-sites assumption may be biased. On the other hand, because the likelihood approach developed here only applies to rare variants, which arose recently, it is not suited to estimate very ancient demographic history.

Thus, because DR EVIL models rare variants, we only infer very recent demography and fix more ancient demography. The ancient demographic history can be fixed by, for example, inferring demographic parameters using infinite-sites site-frequency spectrum methods on a smaller subsample, or using complementary methods, such as coalescent hidden Markov models [68, 69]. We then infer a piecewise constant demography for recent periods. In the Methods, we describe a multi-step approach to infer demographic history and mutation rates jointly.

To test DR EVIL, we simulated data under two different demographic models across sample sizes and compared our estimates with those made using an infinite-sites model. First, we explored a model with two phases of exponential growth (Figure 3). Increasing sample sizes allows for inference of increasingly recent demographic history; with one million samples, we see accurate estimation of the effective population size in the last 10 generations. We also see the critical importance of modeling recurrent mutation: comparing the estimates of effective population size when modeling rare variation (Figure 3A) to estimates that use the infinite-sites model (Figure 3B) shows increased variance and bias in infinite-sites estimates, particularly in the very recent past, where the current effective population size is estimated to be almost three times larger when accounting for recurrent mutation (4,349,954 using infinite sites vs 11,621,304 accounting for recurrent mutation). Supplementary Figure 4 shows a similar pattern under a model of population expansion followed by decrease, in which estimates made using the infinite-sites model are both higher variance and more biased across sample sizes.

**Figure 3:**
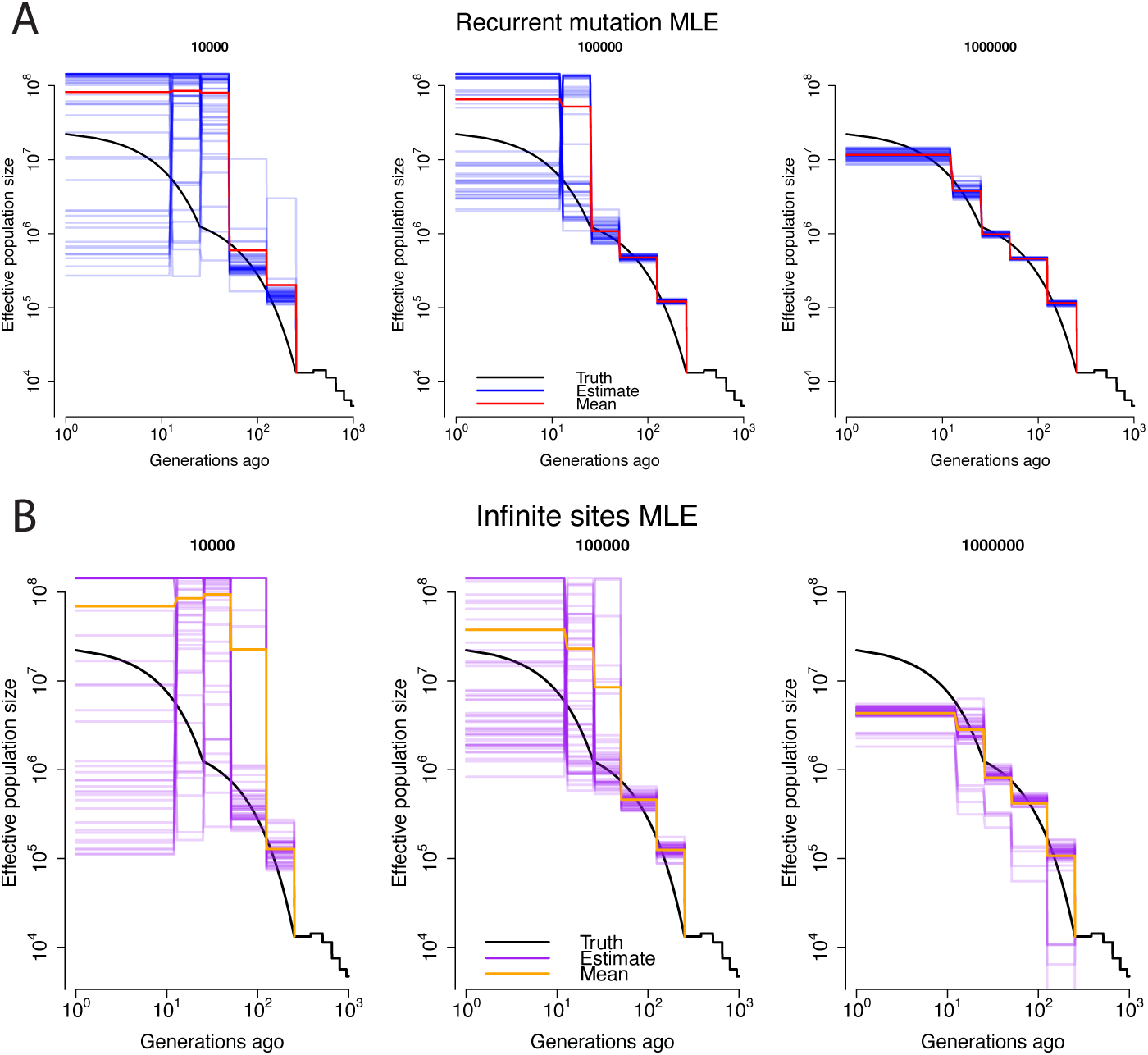
Importance of sample size and recurrent mutation for demographic inference. In each panel, the horizontal axis shows the time in generations and the vertical axis the effective population size. The solid black lines show the simulated demography, while each blue line shows the inference from a single simulation replicate. The red line shows the mean across simulations. The number on top of each sub-panel represents the sample size. A) Estimation accounting for recurrent mutation. B) Estimation under the infinite-sites model.

## 2.3 Application to gnomAD

We applied our approach to estimate mutation rates and recent demographic history from the gnomAD v4.1 dataset. We took advantage of the massive sample size afforded by the inclusion of the UK BioBank exome data and computed the synonymous site-frequency spectrum of non-Finnish European samples, downsampled to 1,000,000 haploids. We imposed an allele-frequency cutoff, only analyzing alleles appearing in the sample fewer than *K* = 1000 times. To account for the context-dependence of mutation rates, we stratified the site-frequency spectrum by trinucleotide context and methylation level. Because some trinucleotide contexts do not result in synonymous mutations, we observed 92 out of the 96 possible trinucleotide contexts. Combined with methylation status at CpG transitions, we analyzed 144 total contexts.

To estimate demographic history, we used the Schiffels–Durbin model estimated using MSMC [69] as a base, and added several more recent time bins (Methods; Supplementary Table 1). We also fit both a single mutation rate and a gamma distribution of mutation rates to each context (Supplementary Table 2). Figure 4A-B shows the fit of the model to the observed data for two trinucleotide contexts, the low-mutation-rate CAA-to-CTA context and the high-mutation-rate CGT-to-CAT context at methylation level 6, respectively. The CAA-to-CTA context appears consistent with an infinite-sites model, but the CGT-to-CAT context can only be fit well by allowing for recurrent mutation. Moreover, we show in Supplementary Figure 5 that the fit is significantly improved by allowing a gamma distribution of mutation rates. Thus, we see that our new model can fit the data well, and that it is important to model recurrent mutation and residual mutation rate variation to do so.

**Figure 4:**
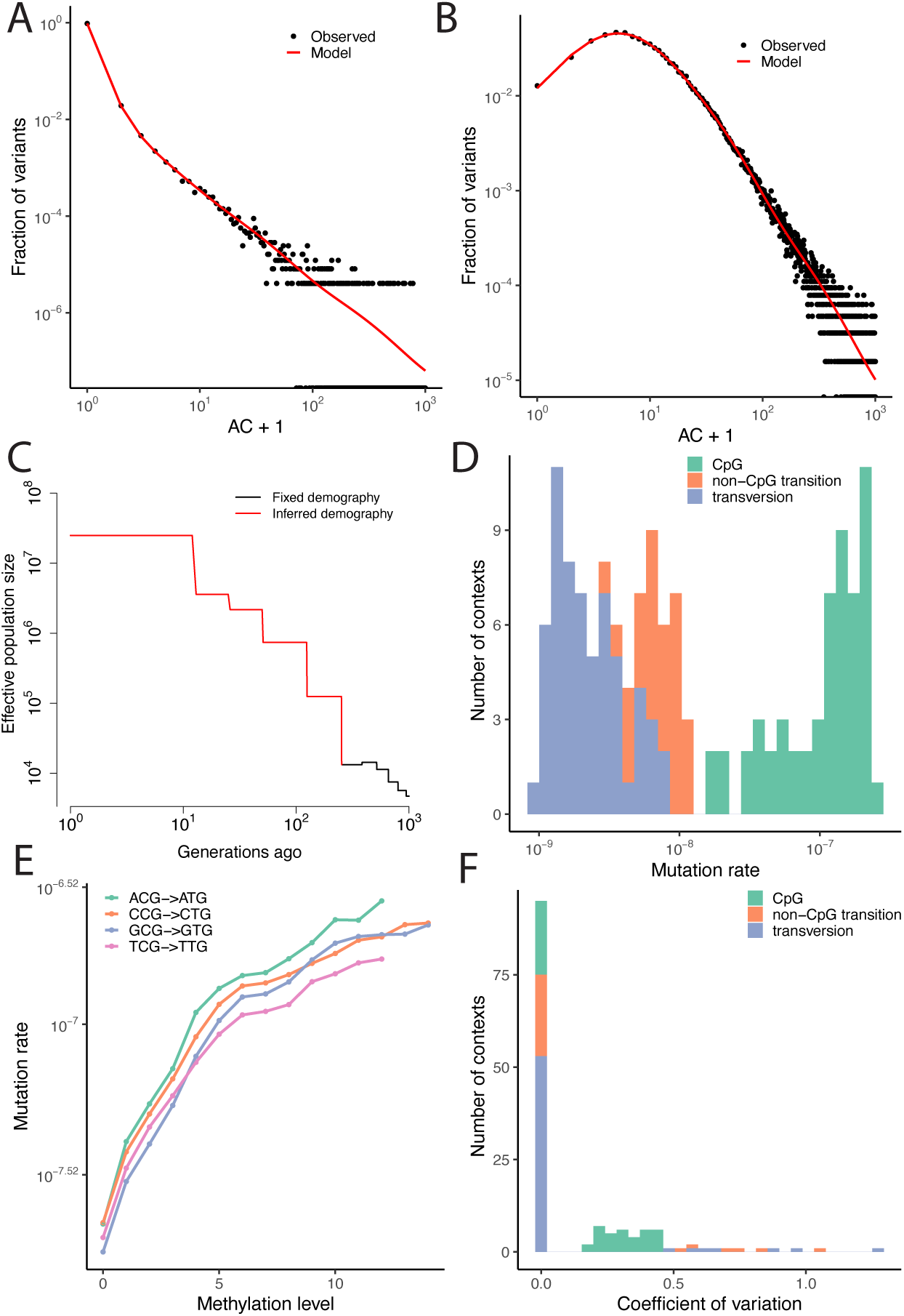
Inference using 1,000,000 samples from gnomAD v4.1. A) The fit of the model for the low-mutation-rate CAA-to-CTA context. B) The fit of the model for the high-mutationrate CGT-to-CAT context at methylation level 6. In panels A and B, the horizontal axis shows the number of minor alleles plus 1, to account for the log scale. C) The estimated demographic history. D) The estimated distribution of mutation rates across contexts and methylation status. E) Mutation rate as a function of methylation level across CpG contexts. F) The estimated coefficients of variation across contexts and methylation status. Contexts in which the null hypothesis of no heterogeneity is not rejected at a 10% FDR are assigned a coefficient of variation of 0.

Figure 4C shows our estimated demographic history. Consistent with expectations, we see explosive population growth and infer an effective population size of approximately 25,000,000 over the last *∼*10 generations. Because the majority of the non-Finnish European cohort in gnomAD v4.1 is from the UK Biobank, this provides an estimate of the UK effective population size that is concordant with a recent estimate of 27 million made using fundamentally different methods [70].

We then examined the distribution of mutation rates we estimated. Figure 4D shows that, consistent with previous analyses, CpG transitions have estimated mutation rates 10 to 100 times larger than other contexts. Supplementary Figure 6 shows that mutation-rate estimates with and without monomorphic sites are consistent for high-mutation-rate contexts, while Supplementary Figure 7 shows that that our estimates are largely concordant with those of gnomAD, although for some contexts we find substantial disagreement.

Because of explosive population growth, all but the lowest mutation-rate categories show substantial deviations from the infinite-sites model, and fits are improved by incorporating recurrent mutation (Supplementary Figure 8). Within CpG contexts, we find that both context and methylation level are associated with mutation-rate differences (Figure 4E). Further, although we fail to reject the null hypothesis of no mutation-rate heterogeneity for most contexts (49/144 contexts have support for heterogeneity at a 10% false discovery rate), Figure 4F shows that we find significant residual mutation-rate variation in many categories, even after accounting for methylation level. Interestingly, while CpG contexts have an appreciable coefficient of variation, our largest estimates of residual variation occur in non-CpG contexts, in line with the observation that there are features beyond methylation and trinucleotide context that influence mutation rates [17, 30].

Finally, we sought to understand the effect of strong selection on variants in large population-variation datasets. Using the demographic model we inferred from gnomAD, we explored the the impact of selection on mutation saturation, allele frequencies, and recurrent mutation. First, we examined the information about selection conveyed by presence or absence of a variant (Figure 5A). Selection affects the ability to observe variants across a range of sample sizes. Unsurprisingly, increasing the sample size results in more observed variants, and increasing the heterozygous selection coefficient results in fewer observed variants. Consistent with previous work [28, 44], positions with high mutation rates are nearly saturated at current sample sizes and essentially completely saturated at samples around 10 million. In contrast, the vast majority of sites with mutation rates between 10*^−^*^9^ and 10*^−^*^8^, will be nonsegregating in samples smaller than 10 million, meaning that presence or absence in a population variation database is only very weakly informative about selection on those mutations.

**Figure 5:**
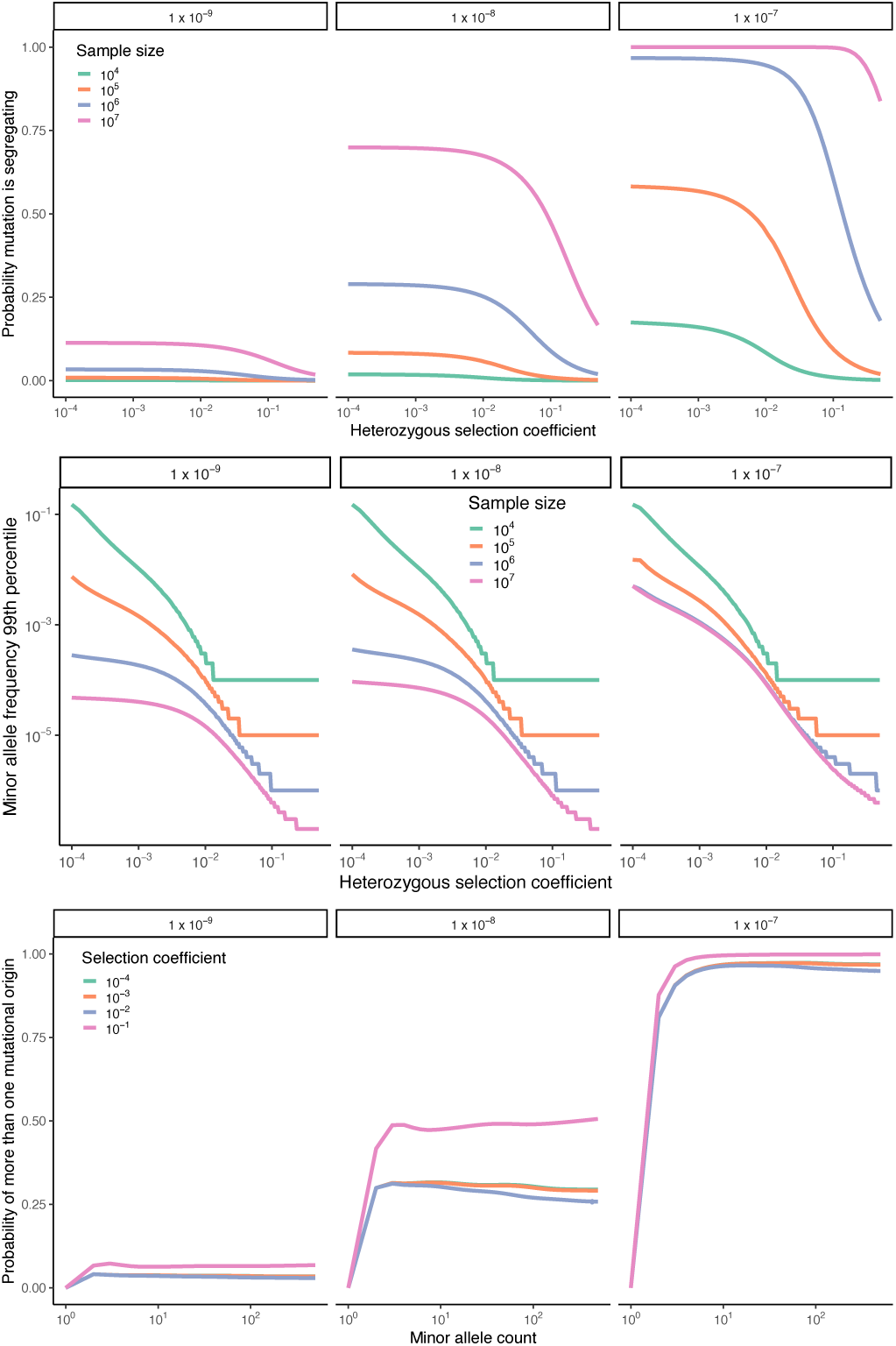
The effect of selection in large samples. A) Strong selection reduces mutational saturation. The horizontal axis shows the heterozygous selection coefficient, while the vertical axis shows the proportion of all possible variants that are observed. Each subpanel corresponds to a different mutation rate, indicated at the top of the panel, and each line corresponds to a sample size, given by the color. B) High mutation rates enable deleterious mutations to rise to high frequency. The horizontal axis shows the heterozygous selection coefficient, and the vertical axis shows the 99th percentile of allele frequencies for segregating alleles. Each subpanel corresponds to a different mutation rate, indicated at the top of the panel, and each line corresponds to a different sample size, indicated by color. C) Mutations have multiple origins in large samples. The horizontal axis shows the minor-allele count in a sample of a million haploid genomes, and the vertical axis shows the probability that a mutation has more than one origin. Each subpanel corresponds to a different mutation rate, indicated at the top of the panel, and each line corresponds to a different heterozygous selection coefficient, indicated by color.

Next, we examined the information available in frequencies of segregating alleles (Figure 5B). At each selection coefficient, we examined the 99th percentile of the allele-frequency distribution conditional on an allele segregating; thus, this is the largest selection coefficient that could be rejected at the 99% level. High mutation rates can result in mutations being an order of magnitude more common at a given selection coefficient compared with lower mutation rates; in samples of size 10 million, CpGs with selection coefficients between 10*^−^*^3^ and 10*^−^*^2^ will be an order of magnitude more common than slower-mutating alleles with the same heterozygous selection coefficient.

We also examined the impact of recurrent mutation (Figure 5C). The prevalence of recurrent mutation has implications for phasing and imputation, which typically assume an infinite-sites model. While alleles with mutation rates on the order of 10*^−^*^9^ are unlikely to have multiple origins, strongly selected mutations with mutation rates on the order of 10*^−^*^8^ or larger have a substantial probability of having arisen multiple times, suggesting that deleterious CpGs may be difficult to phase or impute due to recurrence. As an alternative view of this question, Supplementary Figure 9 shows the expected number of mutational origins, which is close to one for low mutation rates, but can be greater than four for high mutation rates and strong selection. Perhaps surprisingly, we find a non-monotonic relationship between the probability of multiple origins and the selection coefficient, demonstrating the importance of explicit population-genetic modeling to understand how evolutionary forces affect genetic diversity.

### 3 Discussion

Here, we presented DR EVIL, a method for population-genetic estimation of demographic parameters and mutation rates applicable to samples of millions of genomes. Because most of the new information contained in extremely large sequencing studies is contained in the new rare variants they uncover, our approach focuses on modeling rare variation. A focus on rare variation allows computationally efficient handling of recurrent mutation and selection, enabling analyses of large datasets in which the standard neutral infinite-sites assumption is violated.

Our approach allows for estimation of mutation rates with up to an order of magnitude lower root-mean-squared error than standard population-genetic approaches [2, 17]. This is because in large sequencing studies, mutation affects not just the proportion of segregating sites but also the shape of the allele-frequency distribution. Leveraging this fact, we found that in sufficiently large sequencing studies, mutation rates can be estimated without an estimate of the target size; that is, it is unnecessary to account for invariant sites. Computing the number of invariant sites can be difficult: it is important to know whether a site is invariant because it is truly invariant or because it was too difficult to genotype. We also characterized a method-of-moments estimator of the mutation rate that is substantially more robust to recurrent mutation and achieves impressive efficiency at large sample sizes. Further, our method provides a likelihood-ratio test for heterogeneity of mutation rates, and we find evidence of such heterogeneity in many sequence contexts. Our new approach allowed us to make estimates about population history as recently as 10 generations ago, a time period that only appreciably affects the site-frequency spectrum in datasets with hundreds of thousands of haplotypes. Moreover, we found that, because recurrent mutation is a major determinant of allele frequencies for rare variants in ultra-large sequencing studies, traditional population-genetic models that do not account for recurrent mutation result in strongly biased estimates of recent effective population sizes. Our estimate for recent effective population size is concordant with the estimate obtained using identical-by-descent fragments [70]. Thus, our approach provides a “bridge” between SFS-based methods, which excel in the ancient past, and IBD-based methods, which excel in the recent past.

More generally, our work can be seen as broadly in the tradition of site-frequency-spectrum based approaches to demographic estimation [47, 71–73]. Although there are existing theories and methods applicable to samples with two mutations in their history, such as triallelic data [73–75], our method allows for an arbitrary number of mutational origins, so long as the allele remains rare. Multiple mutational origins are expected for high-mutation-rate alleles even in samples of tens of thousands [28, 44]. As we have shown, recurrent mutation is expected to be the norm for rare variants (other than singletons) in high-mutation rate classes given current sample sizes. Our model also adds to the literature on branching-process models of rare variants that allow for recurrent mutation. For neutral variants, our model is identical to that of [54], but we derive an efficient sampling formula and generalize their approach by incorporating natural selection and developing a method for estimation from empirical site-frequency spectra.

Whereas measures of mutational constraint are critical tools to triage genomic regions in disease [76–78], drug discovery [79], and other applications, accurate and interpretable measures of mutational constraint require understanding the demographic and mutational forces that create genetic variation [80]. Using our model, we showed that the interplay between natural selection, mutation, and demography can be complex, supporting the importance of explicit population-genetic modeling for estimating constraint. Moreover, given the possibility of estimation of gene trees from tens or hundreds of thousands of samples [81–84], it may be possible to leverage estimated gene trees to provide enhanced estimates of natural selection. In general, modeling the genealogy of selected variants is difficult [50, 51], but our approach suggests that a fitting a simplified birth-death model may substantially augment our ability to learn about natural selection from gene trees of rare variants.

## 4 Methods

### 4.1 Maximum-likelihood estimation

#### 4.1.1 Likelihood

A key application of sampling formulas in population genetics is to construct a likelihood of the observed data and estimate evolutionary parameters. Here, we are focused on estimation of demographic parameters and mutation rates; we leave inference of natural selection to future work.

The data are the observed allele counts from a sample of haploid size *n*. Because mutation rates vary by genomic context [13, 30] we assume the data are stratified into *J* mutational contexts, where *c_j,k_*is the number of sites of context *j* observed *k* times. We then compute a sampling probability for each context, *p_j,k_*by modifying equation (2) to have a different *θ_j_* for each context, although demographic history is shared across contexts. Finally, because our approximate sampling formula is only valid for rare alleles, we set a maximum allele count *K ≤ n* and renormalize the sampling probability, 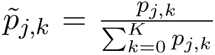
Then, the log-likelihood is

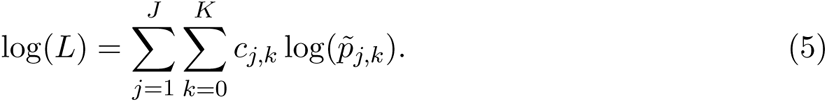

In the Supplementary Material, we prove that this likelihood is valid even when there are more than two alleles segregating at a single locus, as long all alleles but one are rare. Loosely, this is because in the branching-process approximation, the rare alleles do not interfere with each other, just as the different mutational origins of a single rare allele do not interfere with each other.

#### 4.1.2 Estimation procedure

To find maximum-likelihood estimates of mutation rates, we use the fact that the likelihood

1. (5) factorizes by context when the demographic history is fixed. Thus, we can estimate the *θ_c_* for each context independently, which is accomplished by using the one-dimensional Brent optimization algorithm [85] for each context.

To estimate demographic parameters, we begin with the Schiffels–Durbin–Agarwal model [18], which defines a piecewise-constant population size history. We then add two additional pieces, spanning 0–12 generations ago and 12–25 generations ago. We hold the majority of the demographic history fixed and only infer the most recent five pieces of the demographic history, spanning from the present to 252 generations in the past.

To estimate demographic history and mutation jointly, we take a multi-step approach. In the first step, we optimize demographic parameters using the L-BFGS-B algorithm [86] with numerical gradients. During each evaluation of the likelihood, we set *θ_c_* for each context equal to the method-of-moments estimator. This ensures that, given the updated demographic history, the mutation rate parameters are near their maximum-likelihood estimates. Because the likelihood of the site-frequency-spectrum likelihood is known to have complicated geometry [87], we initialize optimization from multiple random locations within demographic parameter space.

Following this initial likelihood optimization, the parameters are near the maximum-likelihood estimates, but not at them, because the method-of-moments estimator of the mutation rate generally differs from the maximum-likelihood estimator. Hence, we perform an additional two rounds of coordinate ascent, in which we alternately fix demographic parameters and find the maximum-likelihood estimators of the mutation rate given the demography, and subsequently fix mutation rates and estimate demographic parameters given the mutation rates.

Finally, we fix the demographic parameters and attempt to fit a gamma distribution of mutation rates to each context. For each context, we compute the likelihood-ratio-test statistic Λ = 2 log(*L*_heterogeneity_*/L*_no heterogeneity_). We assume that under the null hypothesis of no heterogeneity, the likelihood-ratio statistic is distributed as *χ*^2^(1).

### 4.2 Simulations

We developed a custom Wright–Fisher model simulator to simulate unlinked variants evolving subject to genetic drift, natural selection, and recurrent mutation.

To simulate data for testing mutation-rate estimation, we fixed the demographic history at that of the Schiffels–Durbin–Agarwal model [18]. For each mutation rate and sample size, we assumed a target size of 100,000, and imposed a cutoff of *K* = min(0.005 *× n,* 1000) for each sample size *n* for maximum-likelihood analyses, although the method-of-moments and first-order estimators used the entire dataset. We simulated each combination of mutation rate and sample size 45 times and averaged simulations to obtain root mean squared error. When simulating data to test power for inferring mutation-rate heterogeneity, we used the same general scheme with the haploid sample size fixed at one million and simulated mutation rates for each locus from a gamma distribution to generate the desired mean and coefficient of variation. We simulated 40 replicates per combination of mean and coefficient of variation, and computed the likelihood-ratio-test statistic Λ for each. The power estimate is the fraction of tests for which a *χ*^2^ test with one degree of freedom would reject the null hypothesis of no heterogeneity at the 5% level.

To simulate data for estimation of demographic parameters, we began with the Schiffels– Durbin–Agarwal model and modified it to reflect different demographic histories. To simulate two phases of exponential growth, we simulated 227 generations of growth at rate 2% starting 252 generations ago, followed by an additional 25 generations of growth at rate 12%. To simulate a population that expands and contracts, we simulated 227 generations of growth at rate 3% starting 252 generations ago followed by 12 generations of decline at rate *−*10%. For each demographic history, we simulated sample sizes of 1000, 100000, and 1000000. For each demography and sample size pair, we simulated 50 replicates. Each replicate consisted of a simulation of 300000 independently evolving positions, with 100000 having a mutation rate of 10*^−^*^9^ per site per generation, 100000 having a mutation rate of 10*^−^*^8^ per site per generation, and 100000 having a mutation rate of 10*^−^*^7^ per site per generation.

### 4.3 gnomAD data processing

To generate gene annotations, we used the hg38 knownCanonical table from UCSC genome browser [88]. We obtained gene symbols using the kgXref table, and and the exon and coding sequence positions for the canonical transcripts from the knownGene table. We joined tables in the UCSC genome browser by selecting fields from primary and related tables prior to downloading. To ensure that we used only coding exons and not untranslated exons, we restricted only to exons that overlapped with the coding sequence. This resulted in a list of putative genes.

We next used conservation from the 30way phastcons [89] to filter genes. We downloaded the the 30 way phastcons bigwig file from the UCSC genome browser, and used bigWigAverageOverBed to compute the average phastcons score per exon. We then averaged exons together and only retained genes with an average phastcons score greater than

0.3. This resulted in a dataset of 16207 genes. We then computed variant effect predictions (synonymous, nonsynonymous, and stop gain) and trinucleotide contexts for every mutation using a custom script. As a final level of quality control, we joined with gnomAD v4.1 allele number counts, and restricted to sites with a total allele number greater than

1,100,000 that were in the UKBioBank capture region. We also restricted our analyses to only autosomal loci. Any sites that did not pass the quality filters described here were excluded from all subsequent analyses.

We then downloaded variant information from the gnomAD v4.1 dataset from the gnomAD website. For positions without a variant in the gnomAD variant calls, we counted them as having a non-reference frequency of 0. Next, we used a hypergeometric distribution to subsample all sites observed in the non-Finnish European (“nfe”) subset of gnomAD down to a haploid sample size of 1,000,000.

## 5 Software availability

The software to run DR EVIL is available as R code found at https://github.com/ Schraiber/drevil.

## Acknowledgments

This work is dedicated to the memory of Masatoshi Nei. We thank John Wakeley, Louis Fan, Kelley Harris, Alison Feder, Matt Pennell, Molly Przeworski, Guy Sella, Tony Zeng, Hakhamanesh Mostafavi, and Vince Buffalo for helpful discussions during the progress of this work. Funding for JGS and MDE provided by NIH grant R35GM137758.

