## Supplementary Material for "Estimation of demography and mutation rates from one million haploid genomes"

**7 Supplementary Figures** **21**

**8 Supplementary Table Captions** **30**

### 1 Derivation of model

#### 1.1 Sampling from the Wright–Fisher diffusion

We begin with the Wright–Fisher model, which describes the evolution of a variant subject to the combined forces of mutation, genetic drift, and natural selection. We consider the two alleles $A$ (the focal allele, which will later be assumed to be rare) and $a$. For each possible diploid genotype, we define the relative fitnesses $w_{aa} = 1$, $w_{Aa} = 1 + hs$, and $w_{AA} = 1 + s$. We also define mutation from $a$ to $A$ to occur at rate $\mu_1$, while mutation from $A$ to $a$ occurs at rate $\mu_2$. Finally, we suppose that at generation $\tau$, the effective population size is $N_e(\tau)$. We then pass to the diffusion limit, in which case we define new parameters using a reference effective population size, $N_0$. Typically, we take $N_0 \equiv N_e(0)$. Then, we work with compound parameters $t = \frac{\tau}{2N_0}$ $\gamma = 4N_0 s$, $\theta_1 = 4N_0\mu_1$, $\theta_2 = 4N_0\mu_2$, and $\rho(t) = \frac{N_e(2N_0 t)}{N_0}$. Then, the infinitesimal generator of the Wright–Fisher diffusion limit is

$$\mathcal{L}_{WF} = \left(\frac{\gamma}{2}x(1-x)(x+h(1-2x)) + \frac{\theta_1}{2}(1-x) - \frac{\theta_2}{2}x\right)\frac{d}{dx} + \frac{1}{2\rho(t)}x(1-x)\frac{d^2}{dx^2}. \tag{1}$$

We need the probability density that the allele $A$ is at frequency $x$ at time $t$, denoted by $\phi(x;t)$. Standard diffusion theory [1, 2] shows that this density satisfies the partial differential equation

$$\begin{aligned}\frac{\partial}{\partial t}\phi(x;t) &= \mathcal{L}^*_{WF}\phi(x;t)\\ &= -\frac{\partial}{\partial x}\left\{\left(\frac{\gamma}{2}x(1-x)(x+h(1-2x)) + \frac{\theta_1}{2}(1-x) - \frac{\theta_2}{2}x\right)\phi(x;t)\right\}\\ &\quad + \frac{1}{2\rho(t)}\frac{\partial^2}{\partial x^2}\left\{x(1-x)\phi(x;t)\right\},\end{aligned}$$

where $\mathcal{L}^*_{WF}$ is the adjoint operator of $\mathcal{L}_{WF}$.

With a solution for $\phi(x;t)$ in hand, we can model a finite sample of size $n$ taken from the population by binomial sampling. Given the allele frequency $x$, the probability that $k$ out of $n$ alleles are of type $A$ is distributed as a binomial with probability $x$ and size $n$. Averaging over the allele frequency, we see that

$$\begin{aligned}p_{n,k}(t) &= \int_0^1 \binom{n}{k} x^k(1-x)^{n-k}\phi(x;t)dx\\ &= \mathbb{E}_{WF}\left(\binom{n}{k}x^k(1-x)^{n-k}\right),\end{aligned} \tag{2}$$

where $\mathbb{E}_{WF}(\cdot)$ represents the expectation over the diffusion generated by (1). The quantity $p_{n,k}(t)$ represents the *sample frequency spectrum* and there are several tools for computing it, as is calculated in [3–5].

##### 1.2 The rare-variant limit

In this work, we consider the case where the sample size $n$ is very large, and the frequency of the focal ($A$) allele $x$ is very small. In that case, we can make several simplifying approximations. First, following [6] we linearize the diffusion generator around small $x$, discarding the terms $x^2$ and $\theta_i x$. This results in the generator

$$\mathcal{L} = \frac{1}{2}(\gamma h x + \theta_1)\frac{d}{dx} + \frac{1}{2\rho(t)} x \frac{d^2}{dx^2}. \tag{3}$$

In the rest of the manuscript, we abuse notation and simply write $\gamma$ instead of $\gamma h$ and use $\theta$ instead of $\theta_1$, bearing in mind that $\gamma$ represents selection on heterozygotes.

Next, we leverage the fact that $x$ is small and $n$ is large to approximate the binomial distribution in (2) with a Poisson distribution. Then, we have that

$$p_{n,k}(t) = \mathbb{E}\left(\frac{(nx)^k}{k!}e^{-nx}\right), \tag{4}$$

where the expectation is now taken over the diffusion generated by (3). In the rest of the manuscript, we suppress the dependence of $p$ on $n$ for notational simplicity, and refer to $p_k(t) \equiv p_{n,k}(t)$.

To obtain a differential equation for $p_k(t)$, we use Dynkin's formula [1, 2]. This says that the expectation of a function $g(x)$ of a diffusion with generator $\mathcal{G}$ satisfies

$$\frac{d}{dt}\mathbb{E}(g(x)) = \mathbb{E}(\mathcal{G}g(x)).$$

Applying this formula with $g(x)$ given by (4) and the generator (3), we have

$$\frac{d}{dt}\mathbb{E}\left(\frac{(nx)^k}{k!}e^{-nx}\right) = \mathbb{E}\left(\frac{1}{2}(\gamma x + \theta)\frac{d}{dx}\left\{\frac{(nx)^k}{k!}e^{-nx}\right\} + \frac{1}{2\rho(t)} x \frac{d^2}{dx^2}\left\{\frac{(nx)^k}{k!}e^{-nx}\right\}\right).$$

The following steps work through evaluating the derivatives and expectations. We have that

$$\frac{d}{dx}\left\{\frac{(nx)^k}{k!}e^{-nx}\right\} = n\frac{(nx)^{k-1}}{(k-1)!}e^{-nx} - n\frac{(nx)^k}{k!}e^{-nx}$$

and

$$\frac{d^2}{dx^2}\left\{\frac{(nx)^k}{k!}e^{-nx}\right\} = n^2\frac{(nx)^{k-2}}{(k-2)!}e^{-nx} - 2n^2\frac{(nx)^{k-1}}{(k-1)!}e^{-nx} + n^2\frac{(nx)^k}{k!}e^{-nx}.$$

So,

$$\begin{aligned}
\mathbb{E}\left(\frac{1}{2}(\gamma x+\theta)\frac{d}{dx}\left\{\frac{(nx)^k}{k!}e^{-nx}\right\}\right) &= \mathbb{E}\left(\frac{1}{2}(\gamma x+\theta)\left(n\frac{(nx)^{k-1}}{(k-1)!}e^{-nx}-n\frac{(nx)^k}{k!}e^{-nx}\right)\right) \\
&= \frac{\gamma}{2}\left(k\mathbb{E}\left(\frac{(nx)^k}{k!}e^{-nx}\right)-(k+1)\mathbb{E}\left(\frac{(nx)^{k+1}}{(k+1)!}\right)\right) \\
&\quad + \frac{n\theta}{2}\left(\mathbb{E}\left(\frac{(nx)^{k-1}}{(k-1)!}e^{-nx}\right)-\mathbb{E}\left(\frac{(nx)^k}{k!}e^{-nx}\right)\right) \\
&= \frac{\gamma}{2}\left(p_k(t)-p_{k+1}(t)\right)+\frac{n\theta}{2}\left(p_{k-1}(t)-p_k(t)\right).
\end{aligned}$$

The second line follows by linearity of expectation and multiplying terms by 1 in the form e.g. $1=\frac{k}{k}$, and the third line from the definition of $p_k(t)$. Using a similar argument,

$$\begin{aligned}
\mathbb{E}\left(\frac{1}{2\rho(t)}x\frac{d^2}{dx^2}\left\{\frac{(nx)^k}{k!}e^{-nx}\right\}\right) &= \mathbb{E}\left(\frac{1}{2\rho(t)}x\left(n^2\frac{(nx)^{k-2}}{(k-2)!}e^{-nx}-2n^2\frac{(nx)^{k-1}}{(k-1)!}+n^2\frac{(nx)^k}{k!}e^{-nx}\right)\right) \\
&= \frac{n}{2\rho(t)}\left((k-1)\mathbb{E}\left(\frac{(nx)^{k-1}}{(k-1)!}e^{-nx}\right)-2k\mathbb{E}\left(\frac{(nx)^k}{k!}e^{-nx}\right)\right. \\
&\quad \left.+(k+1)\mathbb{E}\left(\frac{(nx)^{k+1}}{(k+1)!}e^{-nx}\right)\right) \\
&= \frac{n}{2\rho(t)}\left((k-1)p_{k-1}(t)-2kp_k(t)+(k+1)p_{k+1}(t)\right).
\end{aligned}$$

Combining these terms and rearranging, we arrive at

$$\frac{d}{dt}p_k(t)=\left(\frac{\theta}{2}n+\frac{n(k-1)}{2\rho(t)}\right)p_{k-1}(t)-\left(\frac{\theta}{2}n+\frac{nk}{\rho(t)}-\frac{\gamma}{2}k\right)p_k(t)+\left(\frac{n(k+1)}{2\rho(t)}-\frac{\gamma(k+1)}{2}\right)p_{k+1}(t), \tag{5}$$

and making the substitutions $b(t)=\frac{n}{2\rho(t)}$, $d(t)=\frac{n}{2\rho(t)}-\frac{\gamma}{2}$, and $f=\frac{\theta}{2}n$, we arrive at the form in the main text.

### 2 Robustness of rare-variant sampling

#### 2.1 Robustness to multiallelic loci

In nucleotide data, up to four alleles can segregate at a locus. Here, we consider a scenario in which one of the nucleotides represents the "major allele" and the the other three are rare "minor alleles". Define $x_i$ to be the frequency of minor allele $i$, $\theta_i$ to be the population-scaled mutation rate from the major allele to minor allele $i$, and $\gamma_i$ to be the population-scaled (heterozygous) selection coefficient on minor allele $i$. Then, in the rare-variant regime, where we assume terms of the form $x_ix_j$ and $\theta_ix_i$ are negligible, the three minor alleles evolve according to a 3-dimensional diffusion with infinitesimal generator

$$\mathcal{L} = \sum_{i=1}^{3} \left( \frac{1}{2}(\gamma_i x_i + \theta_i)\frac{\partial}{\partial x_i} + \frac{1}{2\rho(t)} x_i \frac{\partial^2}{\partial x_i^2} \right). \tag{6}$$

With multiple alleles, the binomial sampling of (2) becomes multinomial sampling. Define $k_i$ to be the observed count of allele $i$, $k = k_1 + k_2 + k_3$ the total count of minor alleles, $k_4 = n - k$ the observed count of the major allele, and $x = x_1 + x_2 + x_3$ be the total frequency of the minor alleles. Under the regime where the sample size becomes large while the frequencies of the three minor alleles stay small, we have that

$$\begin{aligned}
\binom{n}{k_1, k_2, k_3, k_4} x_1^{k_1} x_2^{k_2} x_3^{k_3} x_4^{k_4} &= \frac{n!}{k_1! k_2! k_3! (n-k)!} x_1^{k_1} x_2^{k_2} x_3^{k_3} (1-x)^{n-k} \\
&\sim \frac{\sqrt{2\pi n}\left(\frac{n}{e}\right)^n}{k_1! k_2! k_3! \sqrt{2\pi(n-k)}\left(\frac{n-k}{e}\right)^{n-k}} x_1^{k_1} x_2^{k_2} x_3^{k_3} (1-x)^{n-k} \\
&= \sqrt{\frac{n}{n-k}} \frac{x_1^{k_1}}{k_1!} \frac{x_2^{k_2}}{k_2!} \frac{x_3^{k_3}}{k_3!} \frac{e^{n-k}}{e^n} \frac{n^n}{n^{n-k}} \frac{1}{\left(1-\frac{k}{n}\right)^{n-k}} (1-x)^{n-k} \\
&= \sqrt{\frac{n}{n-k}} \frac{x_1^{k_1}}{k_1!} \frac{x_2^{k_2}}{k_2!} \frac{x_3^{k_3}}{k_3!} e^{-k} n^k \frac{1}{\left(1-\frac{k}{n}\right)^{n-k}} (1-x)^{n-k} \\
&\sim \frac{x_1^{k_1}}{k_1!} \frac{x_2^{k_2}}{k_2!} \frac{x_3^{k_3}}{k_3!} e^{-k} n^k e^k e^{-nx} \\
&= \frac{(nx_1)^{k_1}}{k_1!} e^{nx_1} \frac{(nx_2)^{k_2}}{k_2!} e^{nx_2} \frac{(nx_3)^{k_3}}{k_3!} e^{nx_3},
\end{aligned}$$

i.e. given the population allele frequencies, the observed counts of each minor allele are well approximated as independent draws from Poisson distributions. The second line follows from Stirling's approximation to $n!$, and the third line follows from rewriting $n - k = n(1 - \frac{k}{n})$. The fifth line follows because $\sqrt{\frac{n}{n-k}} \to 1$, $\left(1 - \frac{k}{n}\right)^{n-k} \to e^{-k}$, and $(1-x)^{n-k} = \left(1 - \frac{nx}{n}\right)^{n-k} \to e^{-nx}$ as $n \to \infty$.

Because the observed counts are approximately independent given the allele frequencies, all we need to show is that the allele frequencies are independent, and then we have shown that the observed counts are approximately independent. To prove that the allele frequencies are independent, consider that, by an argument analogous to that for equation 3, the population SFS will be given by a function $\phi(x_1, x_2, x_3)$ that satisfies

$$\frac{\partial}{\partial t}\phi(x_1, x_2, x_3) = \sum_{i=1}^{3} \left( -\frac{1}{2}\frac{\partial}{\partial x_i}\left(\gamma_i x_i + \theta_i\right)\phi(x_1, x_2, x_3) + \frac{1}{2\rho(t)}\frac{\partial^2}{\partial x_i^2} x_i \phi(x_1, x_2, x_3) \right). \tag{7}$$

Here and in the following, we suppress the dependence of $\phi$ on time for notational simplicity.

If the allele frequencies are independent, then they will follow a solution of equation 7 in which $\phi$ factorizes into $\phi(x_1, x_2, x_3) = \phi_1(x_1)\phi_2(x_2)\phi_3(x_3)$. We therefore seek a solution in which we assume that $\phi(x_1, x_2, x_3) = \phi_1(x_1)\phi_2(x_2)\phi_3(x_3)$. Under this assumption,

$$\frac{\partial}{\partial t}\phi_1(x_1)\phi_2(x_2)\phi_3(x_3) = \sum_{i=1}^{3}\left(-\frac{1}{2}\frac{\partial}{\partial x_i}\left(\gamma_i x_i + \theta_i\right)\phi_1(x_1)\phi_2(x_2)\phi_3(x_3) + \frac{1}{2\rho(t)}\frac{\partial^2}{\partial x_i^2}x_i\phi_1(x_1)\phi_2(x_2)\phi_3(x_3)\right).$$

Now, use the product rule on the time derivative

$$\frac{\partial}{\partial t}\phi_1(x_1)\phi_2(x_2)\phi_3(x_3) = \phi_2(x_2)\phi_3(x_3)\frac{\partial}{\partial t}\phi_1(x_1)+\phi_1(x_1)\phi_3(x_3)\frac{\partial}{\partial t}\phi_2(x_2)+\phi_1(x_1)\phi_2(x_2)\frac{\partial}{\partial t}\phi_3(x_3)$$

and note that, for example,

$$\frac{\partial}{\partial x_1}\left(\gamma_1 x_1 + \theta_1\right)\phi_1(x_1)\phi_2(x_2)\phi_3(x_3) = \phi_2(x_2)\phi_3(x_3)\frac{\partial}{\partial x_1}\left(\gamma_1 x_1 + \theta_1\right)\phi_1(x_1)$$

and similarly

$$\frac{\partial^2}{\partial x_1^2}x_i\phi_1(x_1)\phi_2(x_2)\phi_3(x_3) = \phi_2(x_2)\phi_3(x_3)\frac{\partial^2}{\partial x_1^2}x_1\phi_1(x_1).$$

Together, this implies that

$$0 = \sum_{i=1}^{3}\left(\prod_{j\neq i}\phi_j(x_j)\right)\left(\frac{\partial}{\partial t}\phi_i(x_i) - \left(-\frac{1}{2}\frac{\partial}{\partial x_i}\left(\gamma_i x_i + \theta_i\right)\phi_i(x_i) + \frac{1}{2\rho(t)}\frac{\partial^2}{\partial x_i^2}x_i\phi_i(x_i)\right)\right).$$

Dividing both sides by $\phi_1(x_1)\phi_2(x_2)\phi_3(x_3)$, which we can do if we assume that all $\phi_i(x_i) \neq 0$,

$$0 = \sum_{i=1}^{3}\frac{1}{\phi_i(x_i)}\left(\frac{\partial}{\partial t}\phi_i(x_i) - \left(-\frac{1}{2}\frac{\partial}{\partial x_i}\left(\gamma_i x_i + \theta_i\right)\phi_i(x_i) + \frac{1}{2\rho(t)}\frac{\partial^2}{\partial x_i^2}x_i\phi_i(x_i)\right)\right)$$

Now, since each term only depends on one of the $x_i$, each of the three summands must be equal to a constant, as in separation-of-variables arguments for partial differential equations, i.e.

$$\frac{1}{\phi_i(x_i)}\left(\frac{\partial}{\partial t}\phi_i(x_i) - \left(-\frac{1}{2}\frac{\partial}{\partial x_i}\left(\gamma_i x_i + \theta_i\right)\phi_i(x_i) + \frac{1}{2\rho(t)}\frac{\partial^2}{\partial x_i^2}x_i\phi_i(x_i)\right)\right) = c_i.$$

so that

$$\frac{\partial}{\partial t}\phi_i(x_i) = -\frac{1}{2}\frac{\partial}{\partial x_i}\left(\gamma_i x_i + \theta_i\right)\phi_i(x_i) + \frac{1}{2\rho(t)}\frac{\partial^2}{\partial x_i^2}x_i\phi_i(x_i) + c_i\phi_i(x_i).$$

We can use the conservation of probability to see that the constant should be $c_i = 0$, which means that each allele independently follows a branching process diffusion. First note that for each allele, there is a finite probability $\psi_i \equiv \mathbb{P}(x_i = 0)$ that it is at the boundary at 0. Then, marginally, we have that

$$\psi_i + \int_0^\infty \phi_i(x_i)dx_i = 1$$

for all $t$, i.e. that

$$\frac{\partial}{\partial t}\left[\psi_i + \int_0^\infty \phi_i(x_i)dx_i\right] = 0$$

for all $t$.

To determine the differential equation satisfied by the point mass, we require the probabilty flux. The flux in the dimension $x_i$ is given by

$$J_i(x_1, x_2, x_3) = (\gamma_i x_i + \theta_i)\phi(x_1, x_2, x_3) - \frac{1}{2\rho(t)}\frac{\partial}{\partial x_i} x_i \phi(x_1, x_2, x_3)$$

[2, Eq 4.9]. Then, the rate of change at the boundary is the flux at $x_i = 0$ integrated over all possible values of the other variables,

$$\begin{aligned}
\frac{d}{dt}\psi_i &= -\left[\int_{\mathbb{R}_+^2} J_i(x_1, x_2, x_3)dx_j dx_k\right]_{x_i=0} \\
&= -\left[\int_{\mathbb{R}_+^2}\left((\gamma_i x_i + \theta_i)\phi(x_1, x_2, x_3) - \frac{1}{2\rho(t)}\frac{\partial}{\partial x_i} x_i \phi(x_1, x_2, x_3)\right) dx_i dx_j\right]_{x_i=0} \\
&= -\left[(\gamma_i x_i + \theta_i)\phi_i(x_i) - \frac{1}{2\rho(t)}\frac{\partial}{\partial x_i} x_i \phi_i(x_i)\right]_{x_i=0},
\end{aligned}$$

where the final line arises from recognizing that the integral over the other variables simply gives the marginal density in dimension $i$. Hence,

$$\begin{aligned}
\frac{d}{dt}\psi_i &= -\left[(\gamma_i x_i + \theta_i)\phi_i(x_i) - \frac{1}{2\rho(t)}\frac{\partial}{\partial x_i} x_i \phi_i(x_i)\right]_{x_i=0} \\
&= -\left[\frac{1}{2}\left(\gamma_i x + \theta_i\right)\phi_i(x_i) - \frac{1}{2\rho(t)}\phi_i(x_i) - \frac{1}{2\rho(t)} x_i \frac{\partial \phi_i}{\partial x_i}\right]_{x_i=0} \\
&= -\frac{\theta_i}{2}\phi_i(0) + \frac{1}{2\rho(t)}\phi_i(0),
\end{aligned}$$

where the final line comes from requiring $\phi_i(0) < \infty$ and that $\frac{\partial \phi_i}{\partial x_i}|_{x_i=0} < \infty$.

Next, compute the rate of change of the probability density for dimension $i$,

$$\begin{aligned}\frac{\partial}{\partial t}\int_0^\infty \phi_i(x_i)dx_i &= \int_0^\infty \frac{\partial}{\partial t}\phi_i(x_i)dx_i \\ &= \int_0^\infty \left(-\frac{1}{2}\frac{\partial}{\partial x_i}\left(\gamma_i x_i + \theta_i\right)\phi_i(x_i) + \frac{1}{2\rho(t)}\frac{\partial^2}{\partial x_i^2}x_i\phi_i(x_i) + c_i\phi_i(x_i)\right)dx_i\end{aligned}$$

Using linearity of integration and the fundamental theorem of calculus, we work on each term separately. First,

$$\begin{aligned}-\int_0^\infty \frac{1}{2}\frac{\partial}{\partial x_i}\left(\gamma_i x_i + \theta_i\right)\phi_i(x_i)dx_i &= -\frac{1}{2}\left(\gamma_i x_i + \theta_i\right)\phi_i(x_i)|_0^\infty \\ &= -\frac{1}{2}\left(\gamma_i x_i\phi_i(x_i)|_0^\infty + \theta_i\phi_i(x_i)|_0^\infty\right) \\ &= \frac{\theta_i}{2}\phi_i(0),\end{aligned}$$

because we assume $\phi_i(x_i) \to 0$ as $x_i \to \infty$. Also,

$$\begin{aligned}\frac{1}{2\rho(t)}\int_0^\infty \frac{\partial^2}{\partial x_i^2}x_i\phi_i(x_i)dx_i &= \frac{1}{2\rho(t)}\frac{\partial}{\partial x_i}x_i\phi_i(x_i)|_0^\infty \\ &= \frac{1}{2\rho(t)}\left(\phi_i(x_i) + x_i\frac{\partial}{\partial x_i}\phi_i(x_i)\right)\Big|_0^\infty \\ &= -\frac{1}{2\rho(t)}\phi_i(0)\end{aligned}$$

again assuming that $\phi_i(x_i) \to 0$ as $x_i \to \infty$. Finally,

$$\int_0^\infty c_i\phi_i(x_i)dx_i = c_i$$

by the assumption that probability is conserved.

Putting the terms together,

$$\begin{aligned}\frac{\partial}{\partial t}\int_0^\infty \phi_i(x_i)dx_i + \frac{\partial}{\partial t}\psi_i &= \frac{\theta_i}{2}\phi_i(0) - \frac{1}{2\rho(t)}\phi_i(0) - \frac{\theta_i}{2}\phi_i(0) + \frac{1}{2\rho(t)}\phi_i(0) + c_i \\ &= c_i.\end{aligned}$$

However, we require this derivative to be zero, and thus the only choice is $c_i = 0$.

Thus, the sampling probabilities for each allele are independent, and each satisfy the sampling formula in the main text with the appropriate $\theta$ and $\gamma$ terms. Since the original partial differential equation (equation 7) is linear, the constants being identically zero forces all solutions to be of this form, and therefore the solution we identify is unique. This could be shown rigorously using Sturm–Liouville theory, though we do not pursue this here.

#### 3 The solution for the sampling formula

##### 3.1 Explicit formula

The full formula for the solution to the sampling probabilities is

$$p_k(t) = e^{-\xi_0} \frac{B_k(\xi_1, \xi_2, \ldots, \xi_k)}{k!}$$

with

$$\xi_i = i! \int_0^t f q_i(t; s) ds$$

where

$$q_i(t; s) = \begin{cases} 1 - \alpha(t; s) & \text{if } i = 0, \\ (1 - \alpha(t; s))(1 - \beta(t; s))\beta(t; s)^{i-1} & \text{if } i > 0 \end{cases}$$

where

$$\alpha(t; s) = 1 - \frac{e^{R(t;s)}}{W(t; s)}$$

and

$$\beta(t; s) = 1 - \frac{1}{W(t; s)}$$

with

$$W(t; s) = 1 + \int_s^t e^{R(t;u)} b(u) du$$

and finally

$$R(t; s) = \int_s^t (b(u) - d(u)) du.$$

The formulas for calculating $q_i(t; s)$ are essentially identical to those of [7] and based on the classic result of [8], but modified so that the process can start at time $s$ instead of time $t$. Note that $W(t; s)$ has the interpretation of the expected total number of births from a process at time $t$ when started from a single individual at time $s$ (including the birth of that first individual), and $R(t; s)$ is the growth rate from time $s$ to $t$.

##### 3.2 Insights into evolutionary process gained from the analytic solution

Although seemingly complicated, the solution provides detailed insight into the evolutionary processes that shape the history of rare alleles. First, because $\alpha(t; s)$ is the probability that a mutation that arises at time $s$ is lost by time $t$, and mutations arise at rate $f$ per unit time, the quantity $\xi_0 = \int_0^t f(1 - \alpha(t; s)) ds$ gives the expected number of recurrent mutations at a locus. Because mutations occur as a Poisson process, this indicates that the probability of observing zero copies in the sample is the Poisson probability $p_0 = e^{-\xi_0}$, exactly as seen in the solution.

To understand the sampling formula for $k > 0$, recognize that $q_i(t;s)$ is the probability of a mutation starting from one copy and being observed at $i$ copies in the sample, and mutations arise at rate $f$ per unit time; thus, the quantity $\int_0^t f q_i(t;s)ds$ is the expected number of mutations starting from a new mutation that are currently at frequency $i$, corresponding to the entries of the infinite-sites site frequency spectrum. Assuming the conditions that allow the branching-process approximation, each independent mutational origin evolves independently.

Finally, the Bell polynomials encode information about the number of ways to partition a set into subsets. First, define $B_{k,m}(\xi_1, \xi_2, \ldots, \xi_{k-m+1})$ to be the $m$th *partial* Bell polynomial. This polynomial provides information about how to partition a set of size $k$ into $m$ subsets. In particular, each monomial of the form $\xi_{i_1}\xi_{i_2}\cdots\xi_{i_m}$, where $i_1 + i_2 + \ldots + i_m = k$ and some of the $i_j$ can be identical but none are 0, has a coefficient that counts the number of ways to make a set of size $k$ out of $m$ non-empty subsets of size $i_1, i_2, \ldots, i_m$ while accounting for permutations of the elements. Then, the complete Bell polynomial is obtained by adding together the partial Bell polynomials, $B_k(\xi_1, \xi_2, \ldots, \xi_k) = \sum_{m=1}^{k} B_{k,m}(\xi_1, \xi_2, \ldots, \xi_{k-m+1})$ and encodes all possible partitions of a set containing $k$ elements into $1, 2, \ldots, m$ subsets.

Combining these facts, we see that the Bell polynomial term accounts for the different sets of mutational origins that can result in an allele being observed at frequency $k$ in a sample.

The interpretation of the Bell polynomial term is illustrated in Figure 1 for an allele observed at a count of $k = 3$ in a sample. There are three different mutational histories consistent with an allele observed three times in a sample: 1) there is a single mutational origin which is observed three times (represented by $B_{3,1}$); 2) there are two mutational origins, one of which is observed twice, and one of which is observed once (represented by $B_{3,2}$); 3) there are three mutational origins, each of which is observed once (represented by $B_{3,3}$). Note that there is exactly one way to partition a set of size 3 into one subset, three ways to partition a set of size 3 into two subsets, and one way to partition a set of size 3 into 3 subsets, giving the Bell polynomial coefficients of each history, i.e.

$$\begin{aligned} B_3(\xi_1, \xi_2, \xi_3) &= B_{3,1}(\xi_1, \xi_2, \xi_3) + B_{3,2}(\xi_1, \xi_2, \xi_3) + B_{3,3}(\xi_1, \xi_2, \xi_3) \\ &= 1 \cdot \xi_3 + 3 \cdot \xi_1\xi_2 + 1 \cdot \xi_1^3 \end{aligned}$$

where the infinite-sites counts for each mutation reaching the required frequency are multiplied, as under the branching-process approximation, they all evolve independently.

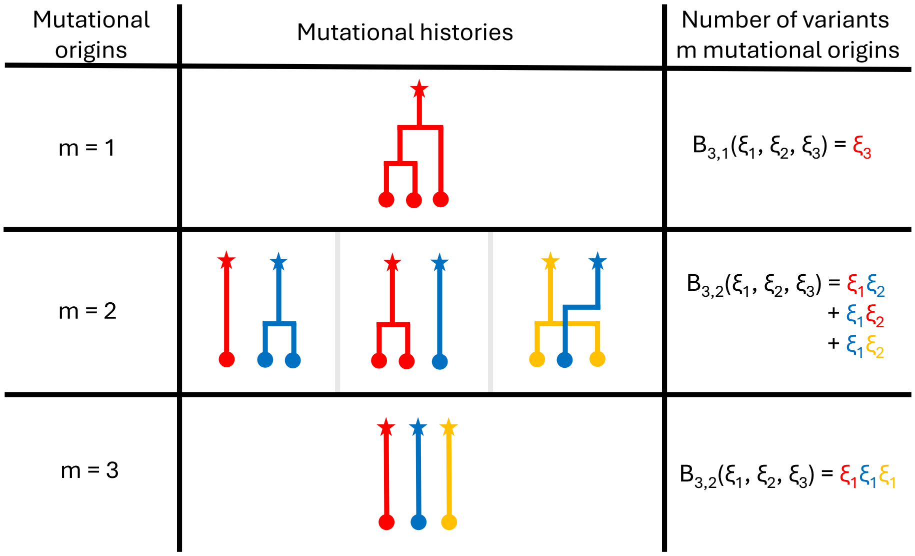

Supplementary Figure 1: Interpretation of the sampling formula. The first column shows the number of mutaitons $m$, and the second column illustrates genealogies for an allele observed $k = 3$ times from $m$ distinct mutations. The final column shows how the partial Bell polynomials count the number of variants that arise from each number of mutational origins.

##### 3.3 Relationship to published formula

The solution in the literature is written (up to different variable names) [9]

$$p_k(t) = \exp\left\{ -\int_0^t \frac{m(u)e^{R(t)-R(u)}}{1+e^{R(t)-R(u)}\left(A(t)-A(u)\right)}du \right\} \frac{B_k(\xi_1, \xi_2, \dots, \xi_k)}{k!}$$

with

$$\xi_i = i! e^{(i-1)R(t)} \int_0^t \frac{f(u)e^{R(t)-R(u)}\left(A(t)-A(u)\right)^{i-1}}{\left(1+e^{R(t)}\left(A(t)-A(u)\right)\right)^{i+1}}du,$$

and

$$A(t) = \int_0^t b(s)e^{-R(s)}ds$$

and

$$R(t) = \int_0^t (b(s) - d(s)ds.$$

Here, we show that our formula can recapitulate this formula. First,

$$\begin{aligned}
1-\alpha(t;s) &= \frac{e^{R(t;s)}}{1+\int_s^t e^{R(t;u)}b(u)du} \\
&= \frac{e^{R(t)-R(s)}}{1+\int_s^t e^{(R(t)-R(u))}b(u)du} \\
&= \frac{e^{R(t)-R(s)}}{1+e^{R(t)}\left(\int_s^t e^{-R(u)}b(u)du\right)} \\
&= \frac{e^{R(t)-R(s)}}{1+e^{R(t)}\left(A(t)-A(s)\right)},
\end{aligned}$$

where, assuming without loss of generality that $t > s > 0$, in the second line we have used

$$\begin{aligned}
R(t;s) &= \int_s^t (b(u)-d(u))du \\
&= \int_0^t (b(u)-d(u))du - \int_0^s (b(u)-d(u))du \\
&= R(t) - R(s)
\end{aligned}$$

and in the fourth line we similarly used

$$\begin{aligned}
\int_s^t e^{-R(u)}b(u)du &= \int_0^t e^{-R(u)}b(u)du - \int_0^s e^{-R(u)}b(u)du \\
&= A(t) - A(s).
\end{aligned}$$

By an almost identical argument,

$$1 - \beta(t;s) = \frac{1}{1 + e^{R(t)}\left(A(t) - A(s)\right)}$$

while

$$\begin{aligned}\beta(t;s) &= 1 - \frac{1}{1 + e^{R(t)}\left(A(t) - A(s)\right)} \\ &= \frac{e^{R(t)}\left(A(t) - A(s)\right)}{1 + e^{R(t)}\left(A(t) - A(s)\right)}\end{aligned}$$

so that

$$\begin{aligned}\xi_i(t;s) &= (1 - \alpha(t;s))(1 - \beta(t;s))\beta(t;s)^{i-1} \\ &= \left(\frac{e^{R(t)-R(s)}}{1 + e^{R(t)}\left(A(t) - A(s)\right)}\right)\left(\frac{1}{1 + e^{R(t)}\left(A(t) - A(s)\right)}\right)\left(\frac{e^{R(t)}\left(A(t) - A(s)\right)}{1 + e^{R(t)}\left(A(t) - A(s)\right)}\right)^{i-1} \\ &= e^{(i-1)R(t)}\frac{e^{R(t)-R(s)}(A(t) - A(s))^{i-1}}{\left(1 + e^{R(t)}\left(A(t) - A(s)\right)\right)^{i+1}}.\end{aligned}$$

Plugging this into the formula for $\xi_i$ in terms of $q_i$ returns the published formula.

### 4 Numerical considerations

#### 4.1 Modified Bell polynomials

The Bell polynomials can be calculated via the recursion

$$B_{k+1}(\xi_1, \xi_2, \ldots, \xi_{k+1}) = \sum_{i=0}^{k} \binom{k}{i} B_{k-i}(\xi_1, \xi_2, \ldots, \xi_{k-i})\xi_{i+1}$$

with initial condition $B_0 = 1$ [10, 11]. To compute a site-frequency spectrum up to an allele count of $K$, we can compute all $K$ Bell polynomials necessary by dynamic programming in $O(K^2)$ time.

Nonetheless, in this formulation, it is not amenable to computation. There is a factor of $i!$ in the formula for $\xi_i$ and the overall formula for $p_k$ is divided by $k!$. Removing the factorial terms is desirable for avoiding overflow in intermediate calculations. To that end, we rewrite the recursion for the Bell polynomials (suppressing the dependence on the $\xi_i$

for convenience)

$$\frac{B_{k+1}}{(k+1)!} = \frac{\sum_{i=0}^{k}\binom{k}{i}B_{k-i}\xi_{i+1}}{(k+1)!} \tag{8}$$
$$= \sum_{i=0}^{k}\frac{k!}{i!(k-i)!}\frac{1}{(k+1)!}B_{k-i}\xi_{i+1}$$
$$= \sum_{i=0}^{k}\frac{i+1}{k+1}\frac{B_{k-i}}{(k-i)!}\frac{\xi_{i+1}}{(i+1)!},$$

where, if we now define $\tilde{B}_k = B_k/k!$ and analogous $\tilde{\xi}_i = \xi_i/i!$, we have

$$\tilde{B}_{k+1} = \sum_{i=0}^{k}\frac{i+1}{k+1}\tilde{B}_{k-i}\tilde{\xi}_{i+1} \tag{9}$$

#### 4.2 Expressions for piecewise-exponential population sizes

When performing calculations, we assume the demographic history is piecewise exponential,

$$\rho(t) = \begin{cases} a_1 e^{\frac{R_1}{2}(t-t_1)}, & \text{if } 0 = t_1 < t \le t_2 \\ a_2 e^{\frac{R_2}{2}(t-t_2)}, & \text{if } t_2 < t \le t_3 \\ \vdots \end{cases}$$

where we have defined the rate $R_i = 4Nr$.

Now, define $I$ to be the largest $i$ such that $t_i < t$, and we have

$$\begin{aligned} A(t) &= \int_0^t \frac{n}{2\rho(s)} e^{-\frac{\gamma}{2}s} ds \\ &= \frac{n}{2}\left(\sum_{i=1}^{I-1}\int_{t_i}^{t_{i+1}} \frac{1}{a_i e^{\frac{R_i}{2}(s-t_i)}} e^{-\frac{\gamma}{2}s} ds + \int_{t_I}^{t} \frac{1}{a_I e^{\frac{R_I}{2}(s-t_I)}} e^{-\frac{\gamma}{2}s} ds\right) \\ &= n\left(\sum_{i=1}^{I-1}\frac{e^{\frac{R_i}{2}t_i}\left(e^{-\frac{\gamma+R_i}{2}t_i} - e^{-\frac{\gamma+R_i}{2}t_{i+1}}\right)}{a_i(\gamma+R_i)} + \frac{e^{\frac{R_I}{2}t_I}\left(e^{-\frac{\gamma+R_I}{2}t_I} - e^{-\frac{\gamma+R_I}{2}t}\right)}{a_I(\gamma+R_I)}\right). \end{aligned}$$

For cases when both $\gamma$ and $R$ are equal to 0, we need to take a limit of the previous formula, arriving at the conclusion that each piece of the integral is

$$\frac{n}{2}\int_{t_i}^{t_{i+1}} \frac{1}{a_i} ds = \frac{n}{2}\frac{t_{i+1}-t_i}{a_i}$$

Now, we intend for $n$ (the sample size of the study) to be large, so we can factor it out of the formula for $A$ derived above. Define $n\tilde{A} = A$, and substitute $R(t) = \frac{\gamma}{2}t$, $f(t) = \frac{\theta}{2}n$. This lets us rewrite

$$\int_0^t \frac{\frac{\theta}{2}ne^{\frac{\gamma}{2}(t-u)}\left(A(t)-A(u)\right)^{i-1}}{\left(1+e^{\frac{\gamma}{2}t}\left(A(t)-A(u)\right)\right)^{i+1}}du = \frac{n}{2}\theta\int_0^t \frac{e^{\frac{\gamma}{2}(t-u)}\left(n\tilde{A}(t)-n\tilde{A}(u)\right)^{i-1}}{\left(1+e^{\gamma t}\left(n\tilde{A}(t)-n\tilde{A}(u)\right)\right)^{i+1}}du$$

$$= \frac{\theta}{2n}\int_0^t \frac{e^{\frac{\gamma}{2}(t-u)}\left(\tilde{A}(t)-\tilde{A}(u)\right)^{i-1}}{\left(\frac{1}{n}+e^{\frac{\gamma}{2}t}\left(\tilde{A}(t)-\tilde{A}(u)\right)\right)^{i+1}}du.$$

Thus, we have the formula

$$\tilde{d_i} = \frac{\theta e^{(i-1)\frac{\gamma}{2}t}}{2n}\int_0^t \frac{e^{\frac{\gamma}{2}(t-u)}\left(\tilde{A}(t)-\tilde{A}(u)\right)^{i-1}}{\left(\frac{1}{n}+e^{\frac{\gamma}{2}t}\left(\tilde{A}(t)-\tilde{A}(u)\right)\right)^{i+1}}du$$

$$= \frac{\theta}{2n}\int_0^t \frac{e^{\frac{\gamma}{2}(it-u)}\left(\tilde{A}(t)-\tilde{A}(u)\right)^{i-1}}{\left(\frac{1}{n}+e^{\frac{\gamma}{2}t}\left(\tilde{A}(t)-\tilde{A}(u)\right)\right)^{i+1}}du$$

where we have taken the exponential term inside the integral to tame some additional overflow issues.

### 5 Derivation of recursion for gamma-distributed mutation rates

Unlike the infinite-sites case, the recurrent mutation model should be sensitive to the whole distribution of mutation rates, not just the mean mutation rate. The formula of the analytic solution suggests a simple approach to analytically integrate over a distribution of mutation rates. Using properties of the Bell polynomials, the solution can be written in the form

$$p_k = e^{-\theta\eta_0}\sum_{m=1}^{k} B_{k,m}(\eta_1, \eta_2, \ldots, \eta_{k-m+1})\theta^i,$$

where the $B_{k,m}$ are the partial Bell polynomials and $\eta_i \equiv \xi_1|_{\theta=0}$. This is because the Bell polynomials are polynomials in the $\xi_i$, which themselves are linear functions of $\theta$. This means that if we can compute exponential moments of $\theta$, then we can perform the integral analytically.

A natural choice to compute exponential moments is to suppose that $\theta \sim \text{Gamma}(\alpha, \beta)$. Denote the distribution of $\theta$ by $f(\theta)$, then we need to compute

$$\begin{aligned}
\int_0^\infty p_k f(\theta) d\theta &= \int_0^\infty e^{-xi_0} \frac{B_k(\xi_1, \xi_2, \ldots, \xi_k)}{k!} f(\theta) d\theta \\
&= \frac{\beta^\alpha}{\Gamma(\alpha)} \int_0^\infty \tilde{B}_k \theta^{\alpha-1} e^{-(\beta+\eta_0)\theta} d\theta \\
&= \frac{\beta^\alpha}{\Gamma(\alpha)} F_k(\alpha, \beta + \eta_0)
\end{aligned}$$

where on the second line we have switched to working with our modified Bell polynomials (8), substituted $\xi_0 = \theta\eta_0$, inserted the probability density of the gamma distribution, and defined

$$F_k(\alpha, \beta) = \int_0^\infty \tilde{B}_k \theta^{\alpha-1} e^{-\beta\theta} d\theta.$$

It will be natural to evaluate this integral using integration by parts and develop a recursive formula. Set

$$\begin{aligned}
& u = \tilde{B}_k \theta^{\alpha-1} \\
\implies & du = ((\alpha - 1)\tilde{B}_k \theta^{\alpha-2} + B'_k \theta^{\alpha-1}) d\theta
\end{aligned}$$

and

$$\begin{aligned}
& dv = e^{-(\beta+\eta_0)\theta} d\theta \\
\implies & v = -\frac{1}{\beta + \eta_0} e^{-(\beta+\eta_0)\theta}.
\end{aligned}$$

First note that $uv$ evaluated at $\theta = 0$ equals 0 because for small $\theta$ it is proportional to $\theta$, while $uv$ evaluated at $\theta = \infty$ is also 0 because the exponential decay term dominates. Thus,

$$\begin{aligned}
\frac{\beta^\alpha}{\Gamma(\alpha)} \int_0^\infty \tilde{B}_k \theta^{\alpha-1} e^{-(\beta+\eta_0)\theta} d\theta = \frac{\beta^\alpha}{\Gamma(\alpha)} \Bigg( & \frac{\alpha - 1}{\beta + \eta_0} \int_0^\infty \tilde{B}_k \theta^{\alpha-2} e^{-(\beta+\eta_0)\theta} d\theta \\
& + \frac{1}{\beta + \eta_0} \int_0^\infty \tilde{B}'_k \theta^{\alpha-1} e^{-(\eta_0+\beta)\theta} d\theta \Bigg).
\end{aligned}$$

To evaluate the first term, use the recursive definition of the Bell polynomials and write

$\tilde{\xi}_i = \theta\tilde{\eta}_i$, where $\tilde{\eta}_i$ does not depend on $\theta$. Then,

$$\begin{aligned}\int_0^\infty \tilde{B}_k \theta^{\alpha-2} e^{-(\beta+\eta_0)\theta} d\theta &= \int_0^\infty \sum_{i=0}^{k-1} \frac{i+1}{k} \tilde{B}_{k-1-i} \tilde{d}_{i+1} \theta^{\alpha-2} e^{-(\beta+\eta_0)\theta} d\theta \\ &= \sum_{i=0}^{k-1} \frac{i+1}{k} \tilde{\eta}_{i+1} \int_0^\infty \tilde{B}_{k-i-1} \theta^{\alpha-1} e^{-(\beta+\eta_0)\theta} d\theta \\ &= \sum_{i=0}^{k-1} \frac{i+1}{k} \tilde{\eta}_{i+1} F_{k-i-1}(\alpha, \beta+\eta_0).\end{aligned}$$

To evaluate the second term, note that the derivative of our modified Bell polynomial with respect to $\theta$ is

$$\frac{d}{d\theta}\tilde{B}_k = \frac{d\tilde{\xi}_1}{d\theta}\frac{d}{d\tilde{\xi}_1}\tilde{B}_k + \ldots + \frac{d\tilde{\xi}_k}{d\theta}\frac{d}{d\tilde{\xi}_k}\tilde{B}_k.$$

Using $\frac{d\tilde{\xi}_i}{d\theta} = \tilde{\eta}_i$, we can compute the derivatives of these modified Bell polynomials using the well known formula for the derivative of the Bell polynomials,

$$\frac{d}{d\xi}B_k = \binom{k}{i} B_{k-i},$$

and the chain rule

$$\begin{aligned}\frac{d}{d\tilde{\xi}_i}\tilde{B}_k &= \frac{d\tilde{B}_k}{dB_k}\frac{dB_k}{d\xi_i}\frac{d\xi_i}{d\tilde{\xi}_i} \\ &= \frac{i!}{k!}\frac{k!}{i!(k-i)!}B_{k-i} \\ &= \frac{B_{k-i}}{(k-i)!} \\ &= \tilde{B}_{k-i}.\end{aligned}$$

Thus,

$$\frac{d}{d\theta}\tilde{B}_k = \sum_{i=1}^{k} B_{k-i}\tilde{\eta}_i.$$

Plugging this in,

$$\begin{aligned}\int_0^\infty \tilde{B}'_k \theta^{\alpha-1} e^{-(\eta_0+\beta)\theta} d\theta &= \int_0^\infty \sum_{i=1}^{k} \tilde{B}_{k-i}\tilde{\eta}_i \theta^{\alpha-1} e^{-(\eta_0+\beta)\theta} d\theta \\ &= \sum_{i=1}^{k} \tilde{\eta}_i \int_0^\infty \tilde{B}_{k-i} \theta^{\alpha-1} e^{-(\eta_0+\beta)\theta} d\theta \\ &= \sum_{i=1}^{k} \tilde{\eta}_i F_{k-i}(\alpha, \beta+\eta_0).\end{aligned}$$

Putting it all together,

$$\begin{aligned}\frac{\beta^\alpha}{\Gamma(\alpha)} F_k(\alpha, \beta+\eta_0) &= \frac{\beta^\alpha}{\Gamma(\alpha)} \int_0^\infty \tilde{B}_k \theta^{\alpha-1} e^{-(\beta+\eta_0)\theta} d\theta \\ &= \frac{\beta^\alpha}{\Gamma(\alpha)} \left( \frac{\alpha-1}{\beta+\eta_0} \int_0^\infty \tilde{B}_k \theta^{\alpha-2} e^{-(\beta+\eta_0)\theta} d\theta \right. \\ &\quad \left. + \frac{1}{\beta+\eta_0} \int_0^\infty \tilde{B}'_k \theta^{\alpha-1} e^{-(\eta_0+\beta)\theta} d\theta \right) \\ &= \frac{\beta^\alpha}{\Gamma(\alpha)} \left( \frac{\alpha-1}{\beta+\eta_0} \sum_{i=0}^{k-1} \frac{i+1}{k} \tilde{\eta}_{i+1} F_{k-i-1}(\alpha, \beta+\eta_0) \right. \\ &\quad \left. + \frac{1}{\beta+\eta_0} \sum_{i=1}^{k} \tilde{\eta}_i F_{k-i}(\alpha, \beta+\eta_0) \right) \\ &= \frac{\beta^\alpha}{\Gamma(\alpha)} \left( \frac{1}{\beta+\eta_0} \sum_{i=1}^{k} \frac{(\alpha-1)i+k}{k} \tilde{\eta}_i F_{k-i}(\alpha, \beta+\eta_0) \right),\end{aligned}$$

where the last line follows by reindexing. This provides us a recursive formula. All we need is the base case,

$$\begin{aligned}F_0(\alpha, \beta) &= \int_0^\infty B_0 \theta^{\alpha-1} e^{-\beta\theta} d\theta \\ &= \int_0^\infty \theta^{\alpha-1} e^{-\beta\theta} d\theta \\ &= \frac{\Gamma(\alpha)}{\beta^\alpha},\end{aligned}$$

and we can analytically integrate over a gamma distribution of mutation rates in $O(K^2)$ time.

### 6 The number of mutational origins

#### 6.1 The distribution of the number of mutational origins

The interpretation of the sampling formula provided in the main text immediately suggests an approach to calculate the posterior probability on the number of latent mutations given a mutation's frequency. The joint probability that an allele is observed $k$ times from $m$ mutational origins is $\mathbb{P}(k, m) = e^{-\xi_0} \frac{B_{k,m}}{k!}$ because the partial Bell polynomial $B_{k,m}$ summarizes the number of ways to partition a set of size $k$ into $m$ subsets. Thus, by Bayes theorem,

$$\begin{aligned}\mathbb{P}(m \mid k) &= \frac{\mathbb{P}(k, m)}{\mathbb{P}(k)} \\ &= \frac{B_{k,m}(\xi_1, \xi_2, \ldots, \xi_{k-m+1})}{B_k(\xi_1, \xi_2, \ldots, \xi_k)}.\end{aligned}$$

A particularly interesting case is the probability that a mutation at frequency $k$ has a single origin. This can be computed as

$$\mathbb{P}(m = 1 \mid k) = \frac{\xi_1}{B_k(\xi_1, \xi_2, \ldots, \xi_k)}.$$

In terms of our modified Bell polynomials (8), this is given by

$$\begin{aligned}\frac{d_k}{B_k} &= \frac{\tilde{\xi}_1 k!}{\tilde{B}_k k!} \\ &= \frac{\tilde{\xi}_k}{\tilde{B}_k}.\end{aligned}$$

The probability of greater than one mutational origin is then given by

$$\mathbb{P}(m > 1 \mid k) = 1 - \frac{\tilde{\xi}_k}{\tilde{B}_k}. \tag{10}$$

#### 6.2 The expected number of mutational origins

The expected number of mutational origins given the observed frequency $k$ can be computed as

$$\mathbb{E}(m \mid k) = \sum_{m=1}^{k} k \frac{B_{k,m}}{B_k}.$$

Here, we develop a different representation of this expectation that is more numerically stable and efficient to compute.

First, the exponential generating function of the partial Bell polynomials is

$$\sum_{k=0}^{\infty}\sum_{m=0}^{k} B_{k,m}\frac{t^k}{k!}u^m = \exp\left\{u\sum_{i=1}^{\infty}\xi_i\frac{t^i}{i!}\right\}.$$

Taking a derivative with respect to $m$ on both sides yields

$$\sum_{k=0}^{\infty}\sum_{m=0}^{k} mB_{k,m}\frac{t^k}{k!}u^{m-1} = \left(\sum_{i=1}^{\infty}\xi_i\frac{t^i}{i!}\right)\exp\left\{u\sum_{i=1}^{\infty}\xi_i\frac{t^i}{i!}\right\}.$$

Evaluating at $u = 1$,

$$\begin{aligned}\sum_{k=0}^{\infty}\sum_{m=0}^{k} mB_{k,m}\frac{t^k}{k!} &= \left(\sum_{i=1}^{\infty}\xi_i\frac{t^i}{i!}\right)\exp\left\{\sum_{i=1}^{\infty}\xi_i\frac{t^i}{i!}\right\}\\ &= \left(\sum_{i=1}^{\infty}\xi_i\frac{t^i}{i!}\right)\left(\sum_{j=0}^{\infty}B_j\frac{t^j}{j!}\right)\\ &= \sum_{i=1}^{\infty}\sum_{j=0}^{\infty}\xi_iB_j\frac{t^{i+j}}{i!j!},\end{aligned}$$

where the second line follows because the exponential generating function of the full Bell polynomials is given by

$$\sum_{j=0}^{\infty}B_j\frac{t^j}{j!} = \exp\left\{\sum_{i=1}^{\infty}\xi_i\frac{t^i}{i!}\right\}.$$

Note that on the left side

$$\sum_{k=0}^{\infty}\sum_{m=0}^{k} mB_{k,m}\frac{t^k}{k!} = \sum_{k=0}^{\infty}\mathbb{E}(m \mid k)\frac{t^k}{k!},$$

so the coefficient of $t^k$ on the right-hand-side will be a representation of the expectation. To find it, note that $j + i = k \implies i = k - j$ and because $B_j = 0$ if $i < 0$ the coefficient is

$$\sum_{i=1}^{k}\frac{\xi_iB_{k-i}}{i!(k-i)!} = \sum_{i=1}^{k}\frac{\xi_iB_{k-i}}{i!(k-i)!}.$$

So,

$$\frac{\mathbb{E}(m \mid k)}{k!} = \sum_{i=1}^{k} \frac{\xi_i B_{k-i}}{i!(k-i)!}$$

$$\implies \mathbb{E}(m \mid k) = \sum_{i=1}^{k} \frac{k!}{i!(k-i)!} \xi_i B_{k-i}$$

$$= \sum_{i=1}^{k} \frac{k}{i} \frac{\binom{k-1}{i-1} B_{k-i}(\xi_1, \xi_2, \ldots, \xi_{k-i}) \xi_i}{B_k(\xi_1, \xi_2, \ldots, \xi_k)}.$$

This formula is enlightening because $\frac{\binom{k-1}{i-1} B_{k-i}(d_1, d_2, \ldots, d_{k-i}) \xi_i}{B_k(\xi_1, \xi_2, \ldots, \xi_k)}$ is the probability that a random allele in the sample of size $k$ comes from a lineage of size $i$ (to see this, note that $\xi_i$ is the "intensity" of lineages of size $i$, we need to chose $k-1$ out of the remaining $n-1$ alleles to be in the same lineage, and then the rest of the lineages need to be distributed as an allele with a frequency of $k-i$). Thus, the expected number of lineages is the allele count $k$ divided by the expectation of one over the lineage size.

In terms of our modified numerically stable Bell polynomials, we have that

$$\sum_{i=1}^{k} \frac{k}{i} \frac{\binom{k-1}{i-1} B_{k-i} \xi_i}{B_k} = \sum_{i=1}^{k} \frac{k}{i} \frac{(k-1)!}{(i-1)!(k-i)!} \frac{\tilde{B}_{k-i}(k-i)! \tilde{\xi}_i i!}{\tilde{B}_k k!}$$

$$= \sum_{i=1}^{k} \frac{\tilde{B}_{k-i} \tilde{\xi}_i}{\tilde{B}_k}.$$

#### 7 Supplementary Figures

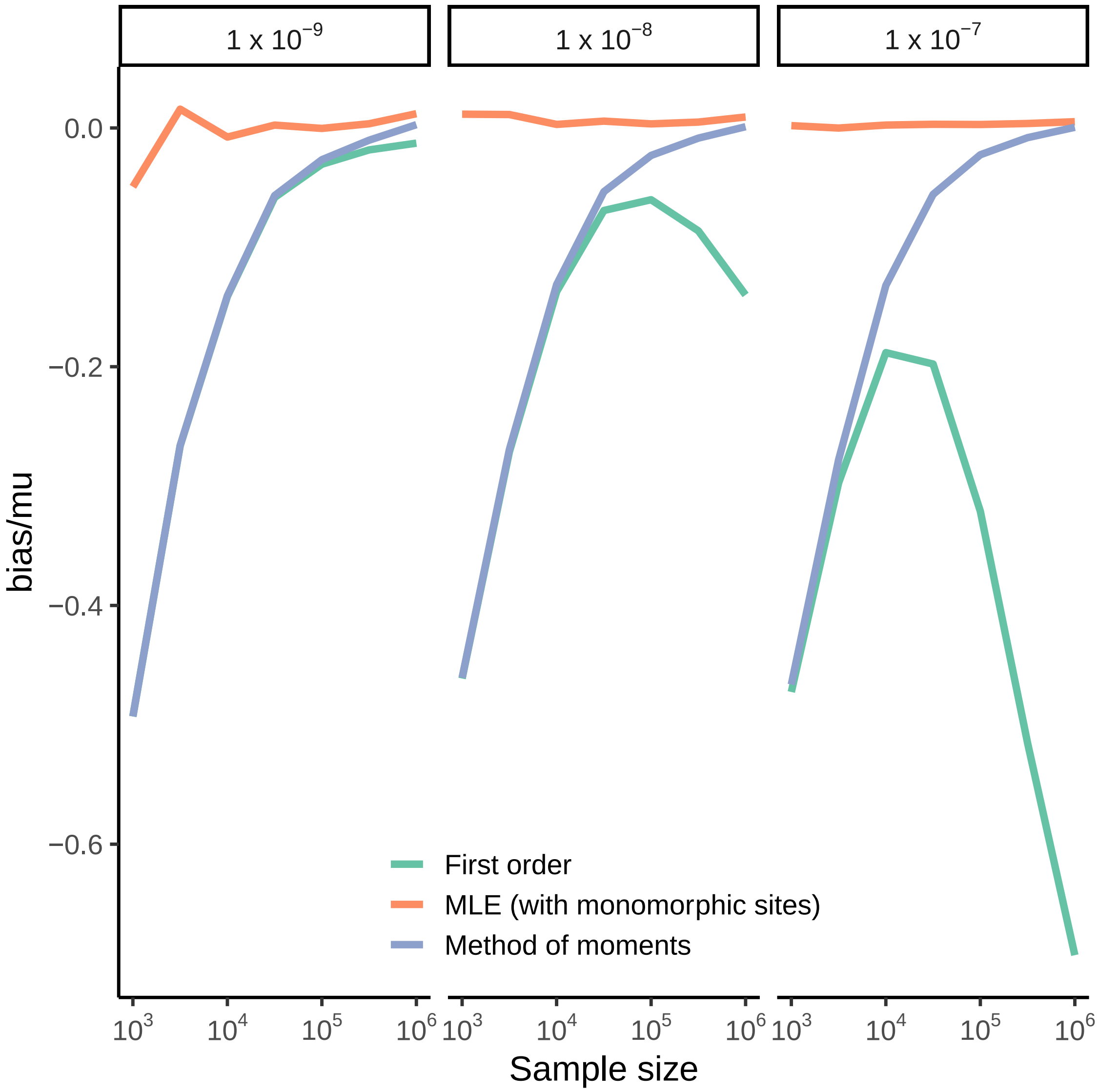

Supplementary Figure 2: The bias of mutation-rate estimators. The horizontal axis shows the sample size and the vertical axis the relative bias. Different methods are indicated with different colors. Each panel shows the results of a simulation with the mutation rate indicated at the top of the panel.

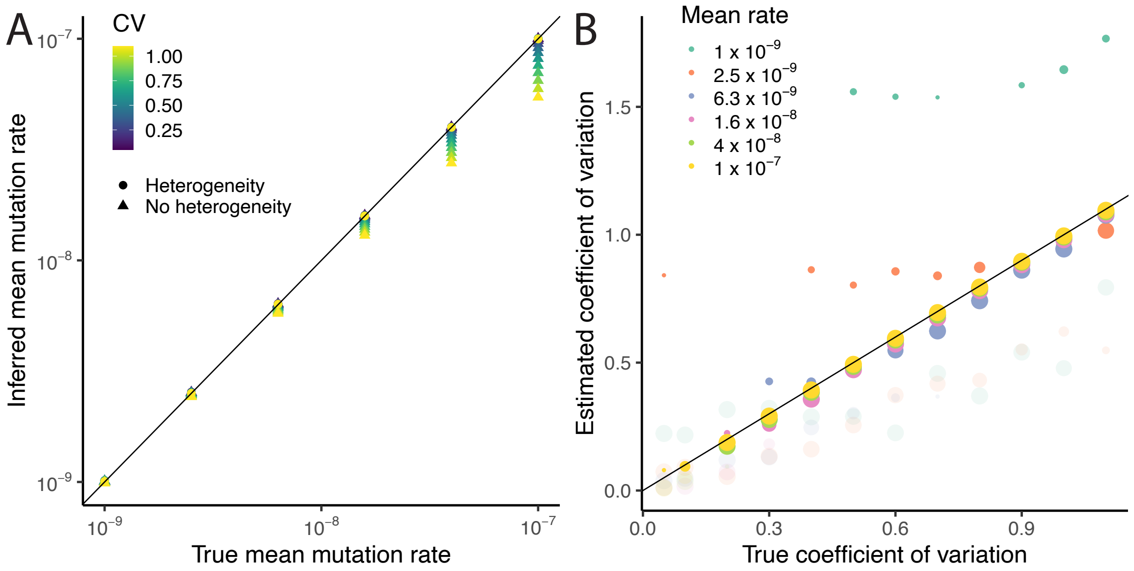

Supplementary Figure 3: Estimation accuracy with mutation-rate heterogeneity. A) Accuracy of mean mutation rate estimation. The horizontal axis shows the mean mutation rate used int he simulations and the vertical axis shows the estimated mutation rate, either when assuming no heterogeneity or heterogeneity, as indicated by the point shape. The point colors indicate the coefficient of variation of the simulation. B) Accuracy of coefficient of variation inference. The horizontal axis shows the coefficient of variation used to simulate data, while the vertical axis shows the maximum-likelihood estimate of the coefficient of variation. The colors of the dots indicate mutation rates, as in the legend. Transparent dots indicate the estimate from simulations that did not reject the null hypothesis of no mutation0rate heterogeneity at the 5%-level. The size of each dot indicates the proportion of simulations at that mutation rate that contribute to each dot.

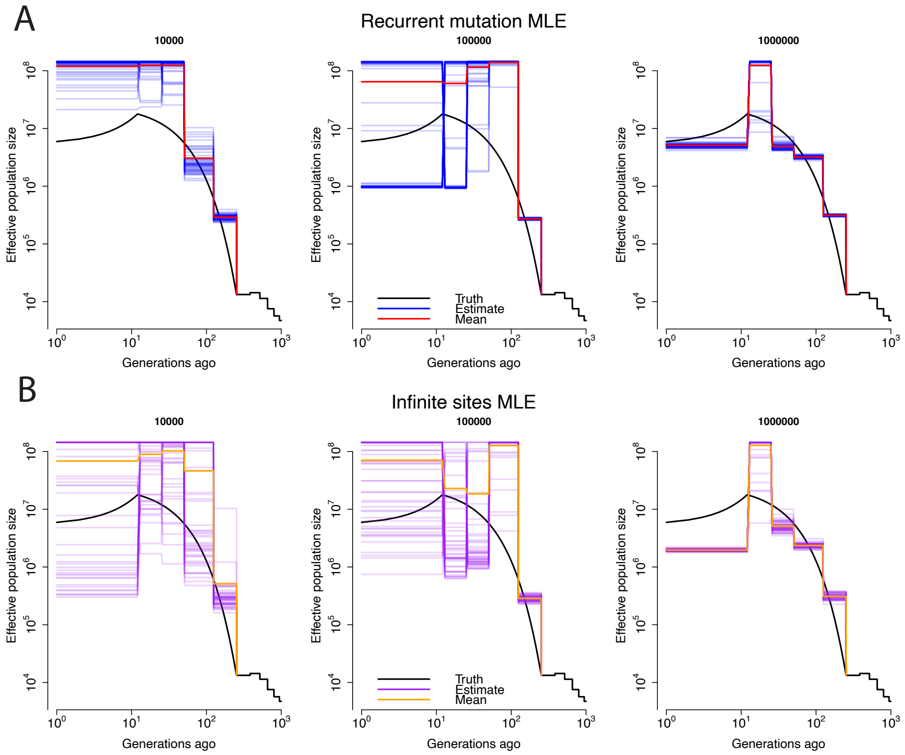

Supplementary Figure 4: Importance of sample size and recurrent mutation for demographic inference under a model of exponential growth followed by exponential decay. In each panel, the horizontal axis shows the time in generations and the vertical axis the effective population size. The solid black lines show the simulated demography, while each blue line shows the estimate from a single simulation replicate. The red line shows the mean across simulations. The number on top of each sub-panel represents the sample size. A) Estimates accounting for recurrent mutation. B) Estimates under the infinite-sites model.

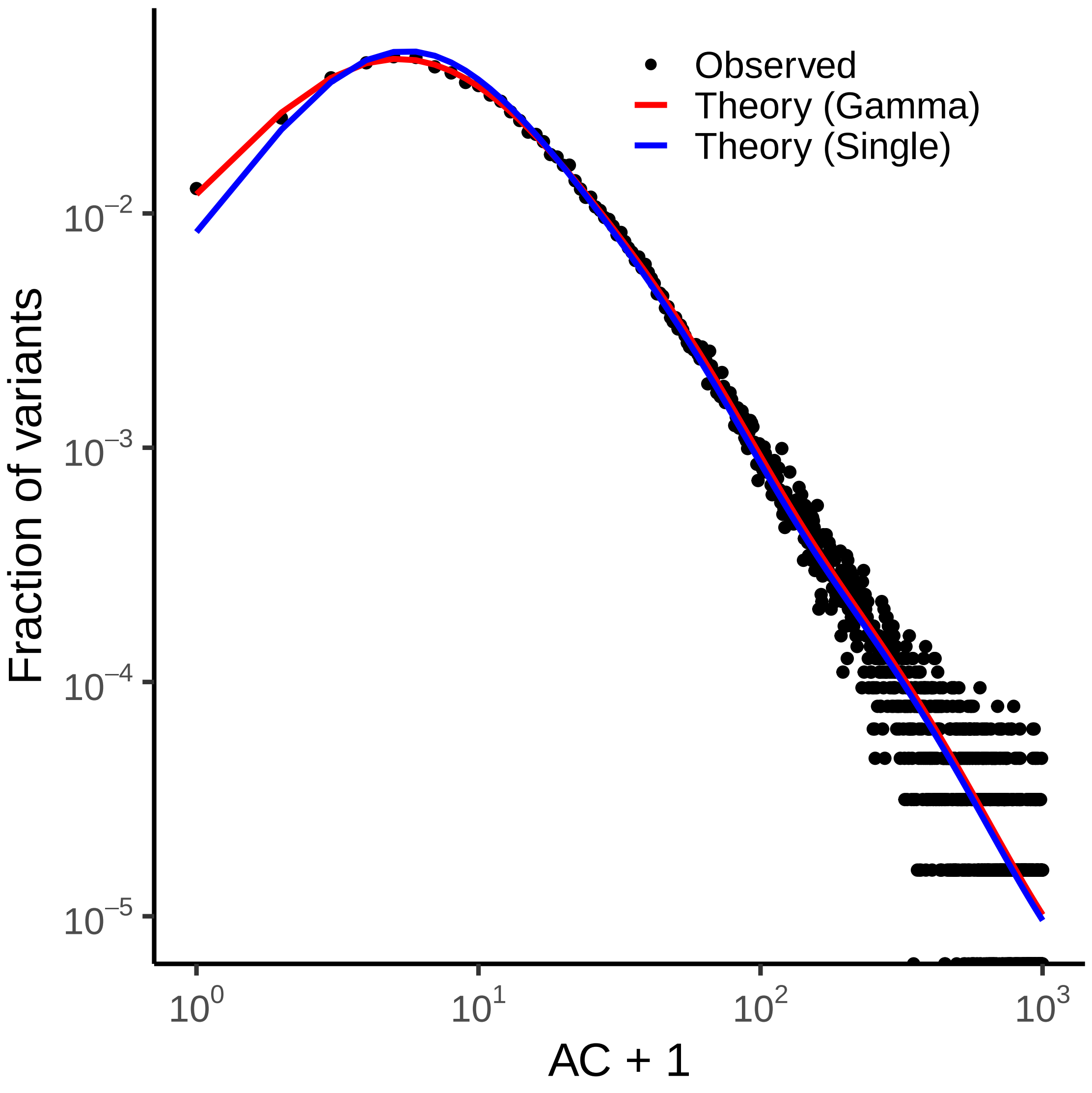

Supplementary Figure 5: The fit to the ACG-to-ATG context with and without mutation-rate heterogeneity. The solid circles show the observed site-frequency spectrum, and the lines indicate the model fit.

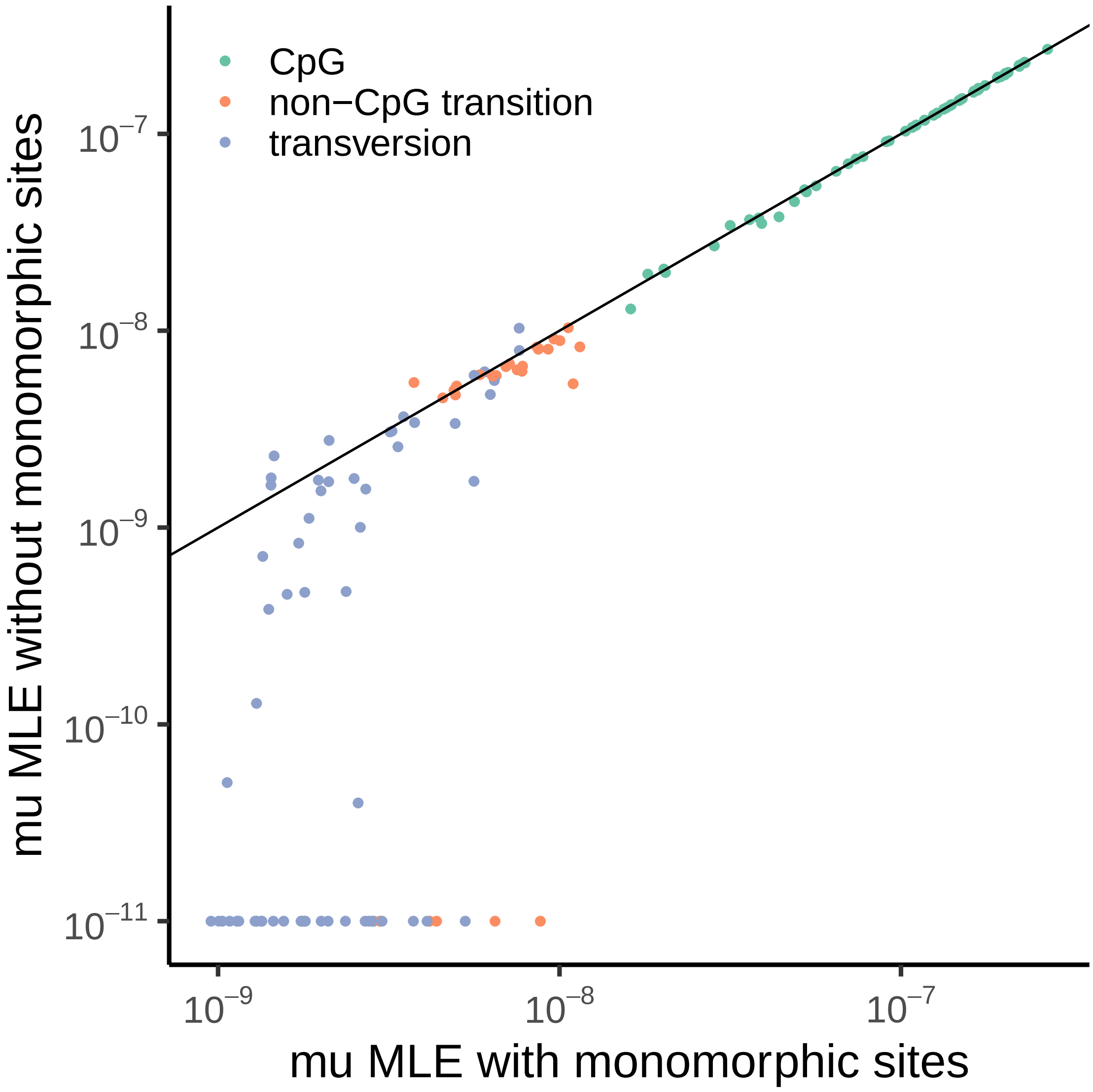

Supplementary Figure 6: Comparison of maximum-likelihood mutation-rate estimates with and without monomorphic sites. The horizontal axis shows the maximum-likelihood estimate of the mutation rate when including sites where no individual has the minor allele, and the vertical axis shows the mutation rate excluding sites where no individual has the minor allele. The line indicates $y = x$, and points lined up at the bottom of the plot have hit the boundary specified on mutation rates. Mutation rates shown are the mean mutation rate of the gamma distribution of mutation rates estimated for each context.

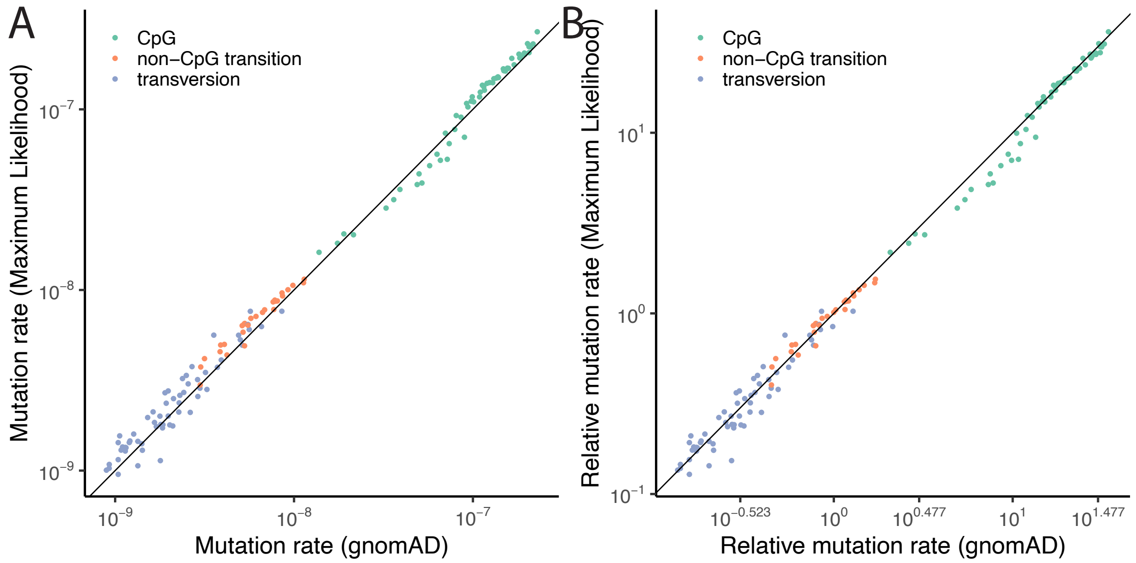

Supplementary Figure 7: Comparison of maximum-likelihood estimates of mutation rates to gnomAD estimates of mutation rates. In each panel, each dot corresponds to a combination of trinucleotide context and methylation level (if applicable). The horizontal axis shows the mutation-rate estimate from gnomAD, while the vertical axis shows the maximum-likeihood mutation rate estimated using our framework. A) Mutation rates are taken directly from our gnomAD and our method. B) Mutation rates are normalized to have mean 1 across variants considered in this study.

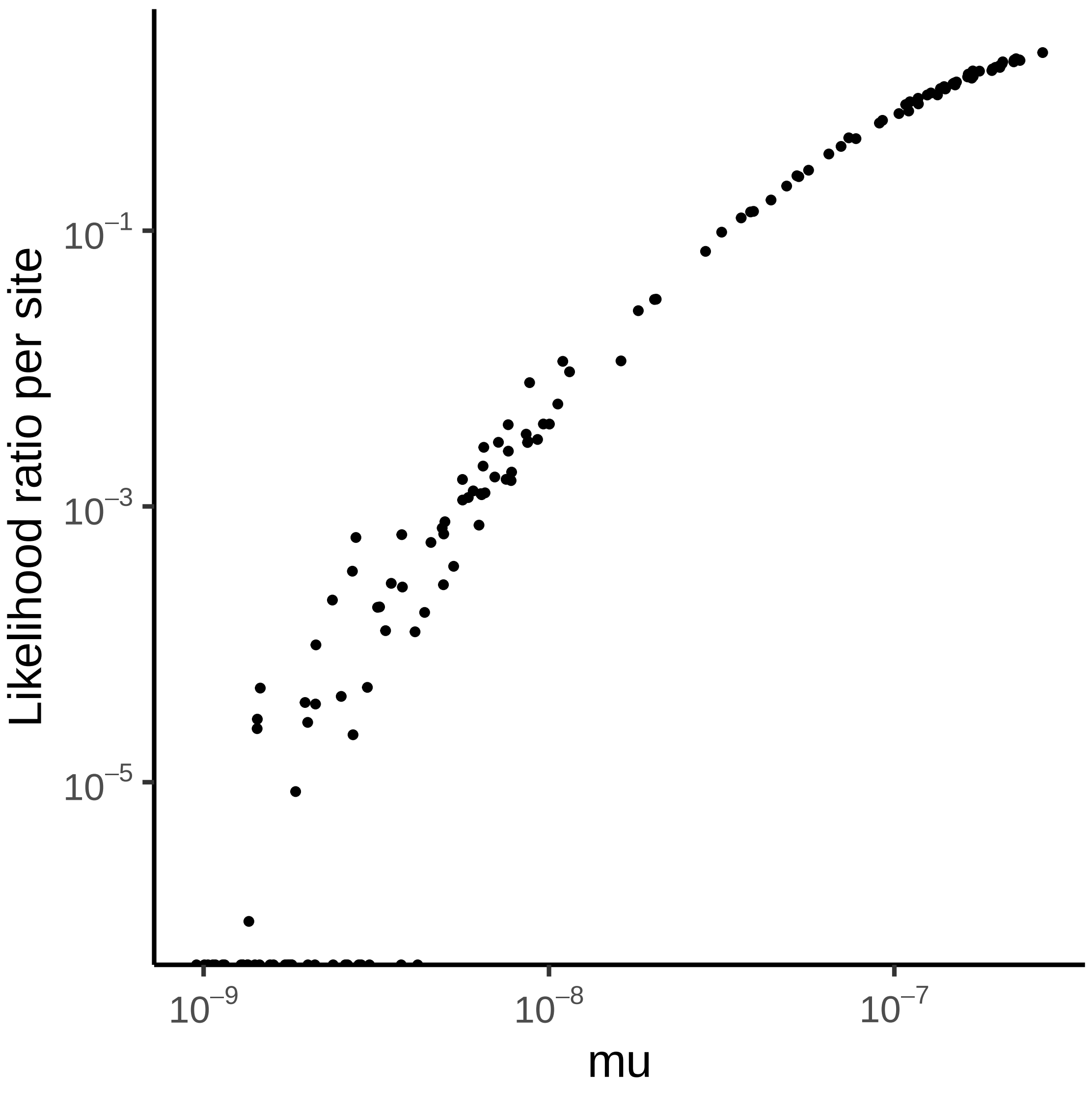

Supplementary Figure 8: Likelihood ratios comparing a recurrent to an infinite-sites model in each sequence context. The horizontal axis shows the estimated mutation rate of the context, and the vertical axis shows the per-site contribution to the likelihood ratio statistic. Computation of the likelihood ratio was only performed on variable sites because monomorphic sites do not make sense under the infinite-sites model. Points at the bottom of the plot indicate a better fit under the infinite-sites model.

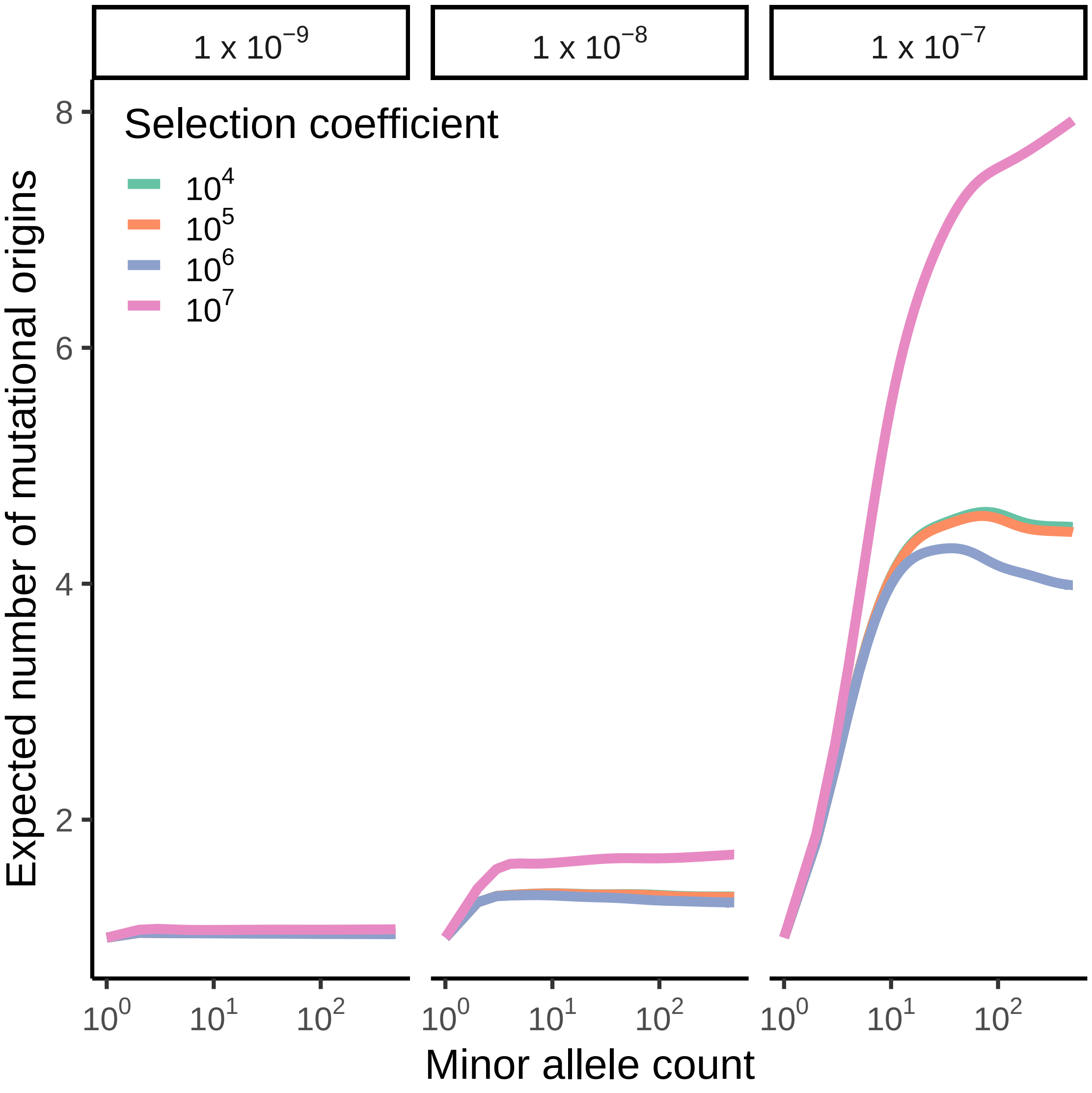

Supplementary Figure 9: The impact of natural selection on the expected number of origins of a mutation. The horizontal axis shows the minor allele count in a sample of size 1000000, and the vertical axis shows the expected number of mutational origins. Each subpanel corresponds to a different mutation rate, indicated at the top of the panel, and each line corresponds to a different selection coefficient, indicated by color.

### 8 Supplementary Table Captions

Supplementary Table 1: Estimated demographic history. The column labeled N_e shows the population size in that piece. The column labeled T_start shows the starting time of that bin in generations. The column labeled inferred indicates whether that Ne was inferred.

Supplementary Table 2: Inferred mutation rates by context.
